# Phospho-signalling deregulation of the mTOR pathway in fibroblasts from LRRK2 Parkinson’s patients

**DOI:** 10.64898/2025.12.05.692089

**Authors:** Alejandro Rubiano-Castro, Joaquín Fernández-Irigoyen, Guillem Parés, Adriana Cortés, Enrique Santamaria, Manel Fernández, Esteban Muñoz, Almudena Sanchez-Gómez, Celia Painous, Ana Cámara, Francesc Valldeoriola, Yaroslau Compta, Eduardo Tolosa, Alicia Garrido, María-José Martí, Rubén Fernández-Santiago, Mario Ezquerra

## Abstract

Activating mutations at the leucine-rich repeat kinase 2 (*LRRK2*) gene are the most frequent genetic cause of Parkinson’s Disease (PD). Using in-depth mass-spectrometry, here we analysed the impact of LRRK2 mutations on the proteome and phospho-proteome of skin fibroblasts from a large LRRK2 clinical cohort from Spain. The cohort encompassed G2019S LRRK2-associated PD (L2PD) (n=15), G2019S LRRK2 non-manifesting carriers (L2NMCs) (n=13), idiopathic PD (iPD) (n=12), and control subjects (n=14) (total n=54). The proteome analysis revealed 89 differential proteins in G2019S L2PD (58 down/ 31 up) and 168 in G2019S L2NMCs (127 down / 41 up) compared to controls. Most of these proteome changes involved deficits in proteins related to mitochondrial energy and ribosomal protein synthesis. At the phospho-proteome level, we identified 394 differential phospho-sites in G2019S L2PD (188 hyper / 206 hypo), 367 in G2019S L2NMCs (215 hyper / 152 hypo), and 428 in iPD (214 hyper / 214 hypo) compared with controls. These phospho-proteomic changes included hyper-phosphorylated proteins associated with the mTOR and Rho GTPase pathways. Notably, the top hyper-phosphorylated hit in G2019S carriers was pThr37/46 4E-BP1, a key effector of the mTOR pathway controlling protein synthesis, and this protein was also deregulated in iPD. In summary, our study uncovers peripheral deregulation of mTOR phospho-signalling associated with LRRK2 mutations, which is related to deficits in mitochondrial and ribosomal proteins, and affects converging pathways altered in iPD.

## INTRODUCTION

Parkinson’s disease (PD) is characterised by the loss of dopaminergic cells in the midbrain substantia nigra and the aggregation of alpha-synuclein across several brain regions, representing the fastest-growing neurodegenerative disorder.^1^ Although most patients present with idiopathic PD (iPD) (95%), pathogenic variants at the leucine-rich repeat kinase 2 (LRRK2) gene causing LRRK2-associated PD (L2PD) are the most common genetic cause of the disease.^2–4^ In addition, LRRK2 non-manifesting carriers (L2NMCs) of pathogenic mutations such as G2019S or R1441G are at an increased risk of PD and constitute a genetic cohort for investigating potential premotor PD stages. At the enzymatic level, LRRK2 variants lead to a toxic gain of kinase function by phosphorylating RAB proteins in disease models,^5^ and endogenous LRRK2 activity biomarkers have been described in blood cells from L2PD patients, including pSer106 RAB12 in G2019S carriers,^6^ and pThr73 RAB10 in R1441G.^7^ Yet, the downstream signalling pathways of mutant LRRK2 in different tissues from patients and the precise pathophysiological mechanisms triggering the expressivity of LRRK2 mutations remain unknown.

Cumulative evidence supports the involvement of peripheral tissues beyond the central nervous system (CNS) in PD, even at prodromal PD stages.^6,8^ In this context, primary skin fibroblasts represent an easily accessible biomatrix which is suitable to investigate the phospho-signalling cascade of mutant LRRK2 in clinical cohorts.^9^ PD fibroblasts have shown alterations associated with PD at multiple levels, including the transcriptome,^10,11^ proteome,^12^ vesicle trafficking,^13^ and mitochondria.^14,15^ Most of these processes have also been reported using induced pluripotent stem cells (iPSC)-derived dopaminergic neurons generated upon reprogramming of LRRK2 patient fibroblasts.^16^ Here, or the first time, we screened the phospho-/proteome of mutant LRRK2 by data-independent acquisition (DIA) mass-spectrometry (MS) using human fibroblasts from a large LRRK2 clinical cohort from Spain, which included G2019S L2PD patients (n=15), G2019S L2NMCs (n=13), iPD cases (n=12), and controls (n=14) (Total n=54) (**Fig. 1**). The main goal of our study was to characterise endogenous phospho-/proteomic alterations in peripheral tissues from LRRK2 mutation carriers, with and without motor symptoms, and to investigate common alterations with iPD.

**Figure 1.**
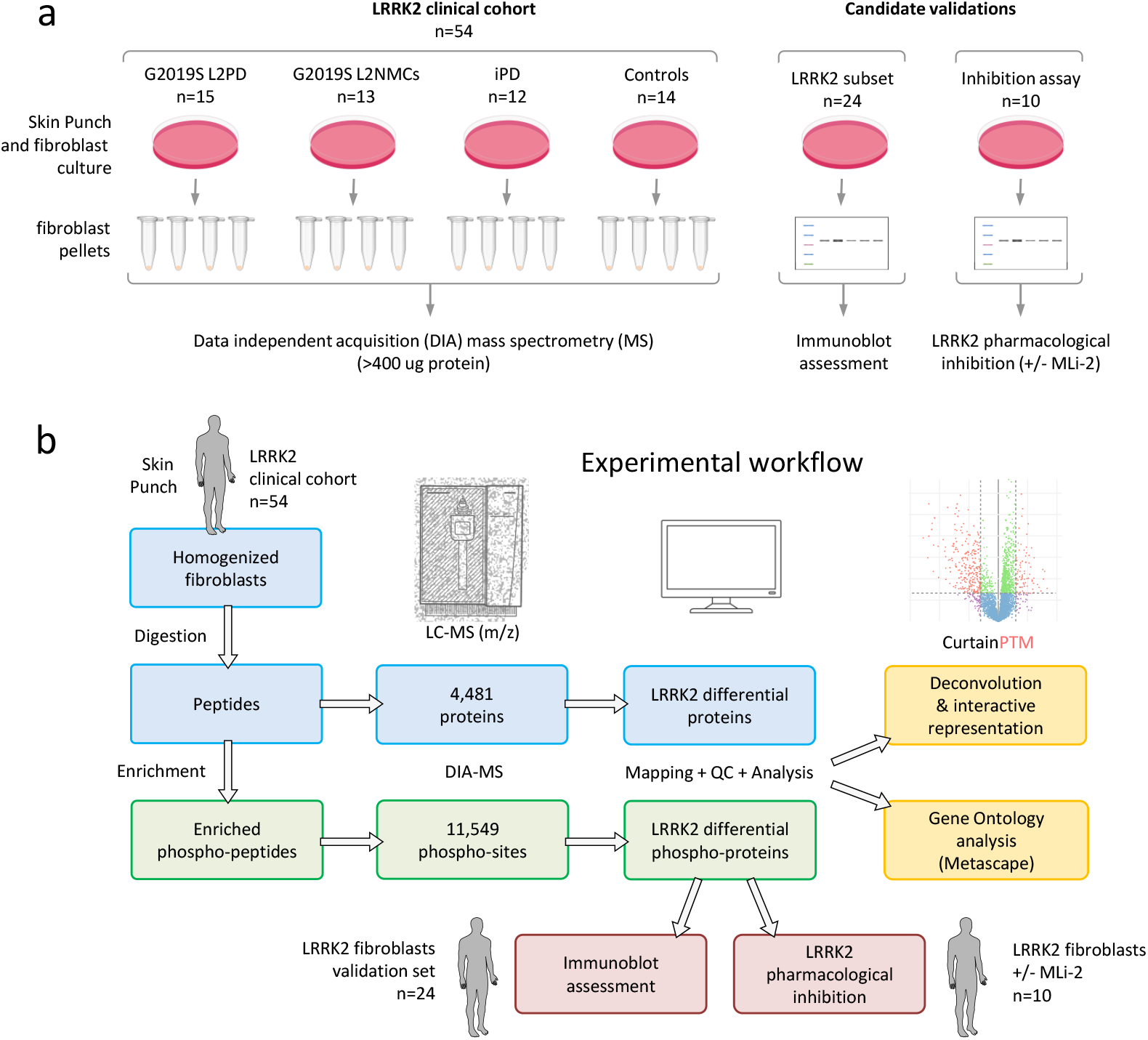
Experimental workflow using fibroblasts from an LRRK2 cohort. **a)** Primary skin biopsies were collected from subjects of an LRRK2 clinical cohort from Spain (n=54) encompassing G2019S L2PD patients (n=15), G2019S L2NMCs (n=13), iPD (n=12), and healthy controls (n=14). **b)** After fibroblast harvesting, homogenisation, and protein digestion, DIA-MS identified 4,481 proteins on an EZ-Exploris 480 mass-spectrometer, and 11,549 phospho-sites after phospho-enrichment. For the group differential analysis, we used a significance cut-off of log_2_FC > 0.6 and a P < 0.05. An interactive representation of the data was done using the Curtain PTM Tool, and Gene Ontology was performed using Metascape. By immunoblot, we assessed pThr37/46 4E-BP1, total ATG9 and total MRPS14 levels in fibroblasts from a subset of subjects (n=24), including G2019S L2PD (n=6), G2019S L2NMCs (n=6), iPD (n=6), and healthy controls (n=6). Lastly, in a second subset of subjects (n=10) encompassing G2019S L2PD (n=4), G2019S L2NMCs (n=2), iPD (n=2) and healthy controls (n=2), treated with DMSO or the MLi-2 LRRK2 inhibitor, we performed an LRRK2 kinase inhibition assay measuring pThr37/46 4E-BP1, pSer106 RAB12, pThr73 RAB10 and pSer935 LRRK2 levels.

## RESULTS

### Proteome analysis of LRRK2 fibroblasts

By in-depth mass-spectrometry, we identified a total of 4,481 proteins expressed in human fibroblasts. Using pairwise comparisons, under a cut-off of log_2_FC>|0.60| and a P<0.05, we identified 89 differential proteins (58 down / 31 up) in L2PD patients vs controls, 65% of which were down-regulated hits, whereas L2NMCs vs controls had 168 differential proteins (127 down / 41 up), involving 76% down-regulated hits. A total of 47% (42 proteins) of the proteome changes from L2PD were shared with L2NMCs, 99% of which had the same fold-change direction. Indeed, L2PD vs L2NMCs revealed the lowest level of proteome changes among all comparisons, with only 76 differential hits (49 down / 27 up), 64% of which were down-regulated in L2PD patients (**Fig. 2; Supplementary Data 1**). In addition, iPD vs controls revealed 358 differential proteins (270 down / 88 up), representing the highest level of proteome differences in the study, with 75% of these hits, similarly to G2019S carriers, being down-regulated proteins in iPD cases. Lastly, 69% of the hits in L2PD (61 proteins) were also common to iPD, indicating a substantial proteome overlap between both groups of PD patients. Altogether, these findings indicate proteome alterations in symptomatic and asymptomatic LRRK2 G2019S carriers involving mostly protein deficits, many of which were down-regulated proteins in iPD as well.

**Figure 2.**
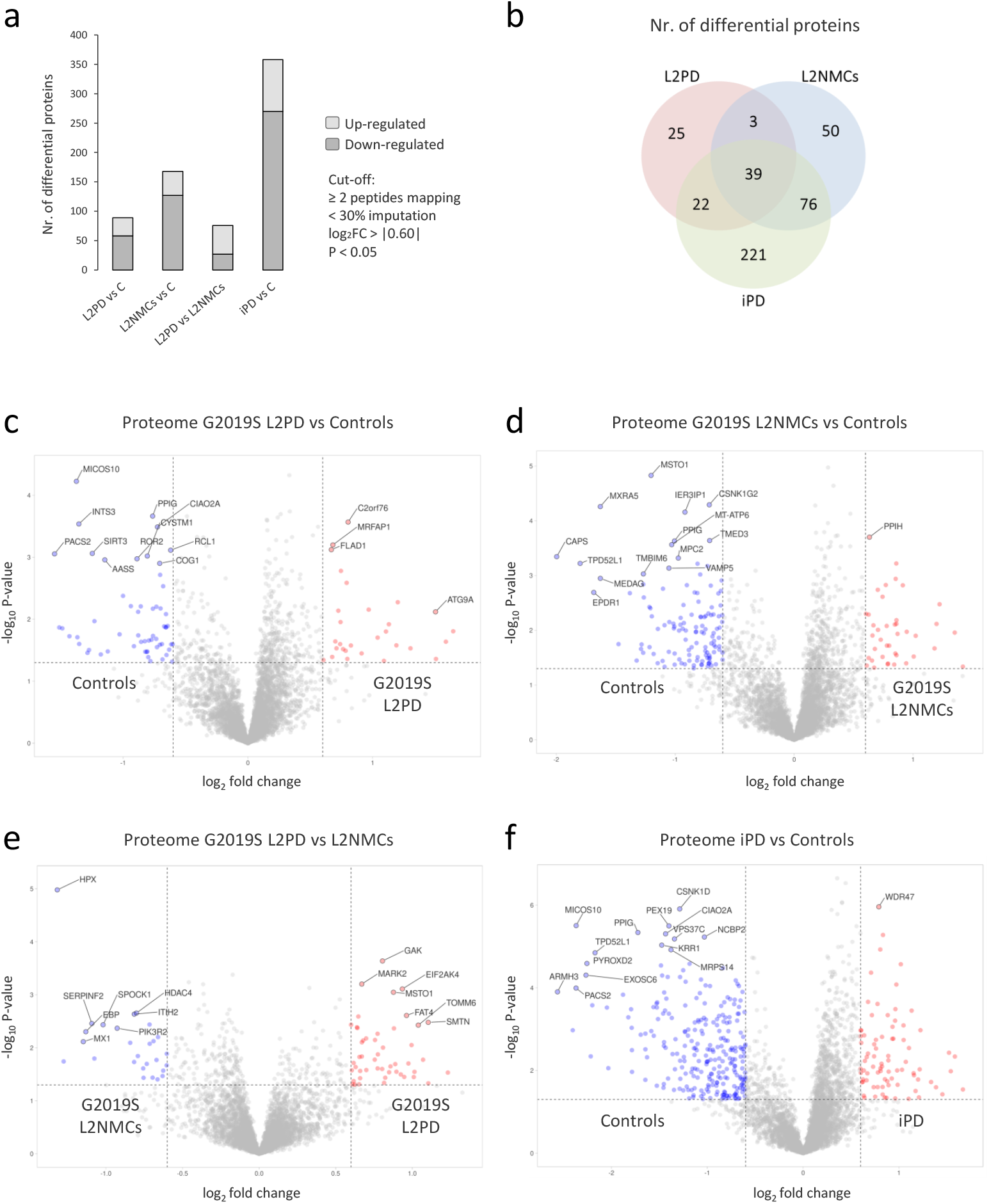
Proteome overview and differential analyses. **a)** Bar plot showing the number of differential proteins in different pairwise comparisons, with up-regulated proteins in dark grey and down-regulated in light grey. The significance cut-off was set at log_2_FC≥⍰0.60⍰ and P<0.05. **b)** Venn diagram showing shared proteins between G2019S L2PD, G2019S L2NMCs, and iPD comparisons vs controls. **c)** Volcano plots showing protein expression differences of the G2019S L2PD group compared to controls. Differential proteins are denoted in red (up-regulated) and blue (down-regulated). **d)** Proteome outcomes of the G2019S L2NMCs compared to controls. **e)** Proteome outcomes of the iPD group compared to controls. **f)** Proteome outcomes of the G2019S L2PD group compared to G2019S L2NMCs. Volcano plots were generated using default parameters in the VolcaNoSer software. Annotated dots correspond to the fifteen protein data points with the largest (Manhattan) distance from the origin above the significance thresholds indicated by the dashed line. VolcaNoSer: https://huygens.science.uva.nl/VolcaNoseR2/.

### LRRK2 proteome top differential hits

Focusing on top individual hits, we observed that the strongest protein hits common to G2019S L2PD, L2NMCs and iPD were down-regulated proteins associated with mitochondrial and ribosomal functions (**Fig. 2**). Among these, protein deficits from L2PD that were common to iPD included MICOS10, which plays a crucial role in maintaining the structure of the mitochondrial membrane;^17^ PACS2, which is involved in mitochondrion-endoplasmic reticulum membrane tethering;^18^ or SIRT3, which is a mitochondrial deacetylase.^19^ Protein deficits in G2019S L2NMCs included MSTO1, related to mitophagy function and fusion,^20^ and MT-ATP6, a component of mitochondrial ATP synthase Complex V.^21^ Lastly, other protein deficits in iPD encompassed MRPS14, which is a protein associated with mitochondrial ribosomes.^22^ In contrast, some differential proteins were prominently up-regulated in both L2PD and iPD, such as ATG9A, which is involved in endosomal processing^12^ and mitochondria autophagy,^23^. In summary, the most significantly deregulated proteins in G2019S mutation carriers and iPD patients included mostly strong protein deficits, which are associated with mitochondrial function and proteostasis.

### Phospho-proteome analysis of LRRK2 fibroblasts

At the phospho-proteome level, we identified 11,549 phospho-sites from 2,732 unique proteins in human fibroblasts, given that individual proteins can contain multiple phosphorylation sites. By pairwise comparison and using the same cut-off as above, we identified 394 unique differential phospho-peptides (188 hyper / 206 hypo) in G2019S L2PD patients, whereas G2019S L2NMCs vs controls had 367 unique differential phospho-peptides (215 hyper / 152 hypo). The overlap of phospho-peptide changes in L2PD and L2NMCs was of only 18% (63 hits), yet 99% of these exhibited full concordant fold-change directions. Remarkably, G2019S L2PD patients compared with G2019S L2NMCs revealed the largest number of phosphorylation changes in the entire study, with 485 differential phospho-peptides (279 hyper / 206 hypo) (**Fig. 3**; **Supplementary Data 2**), suggesting substantial differences related to changes of disease status between G2019S carriers. Lastly, iPD vs controls had 428 differential phospho-proteins (214 hyper / 214 hypo). In addition, only 15% (58 hits) of the differential hits from iPD patients overlapped with L2PD, yet all with full fold-change coincidence. In summary, we observed large phospho-proteome changes in G2019S carriers, with and without motor symptoms, as well as in iPD cases, with highly concordant change directions, and identified shared phosphorylation deregulation affecting the LRRK2 pathway, which is central to the disease pathophysiology.

**Figure 3.**
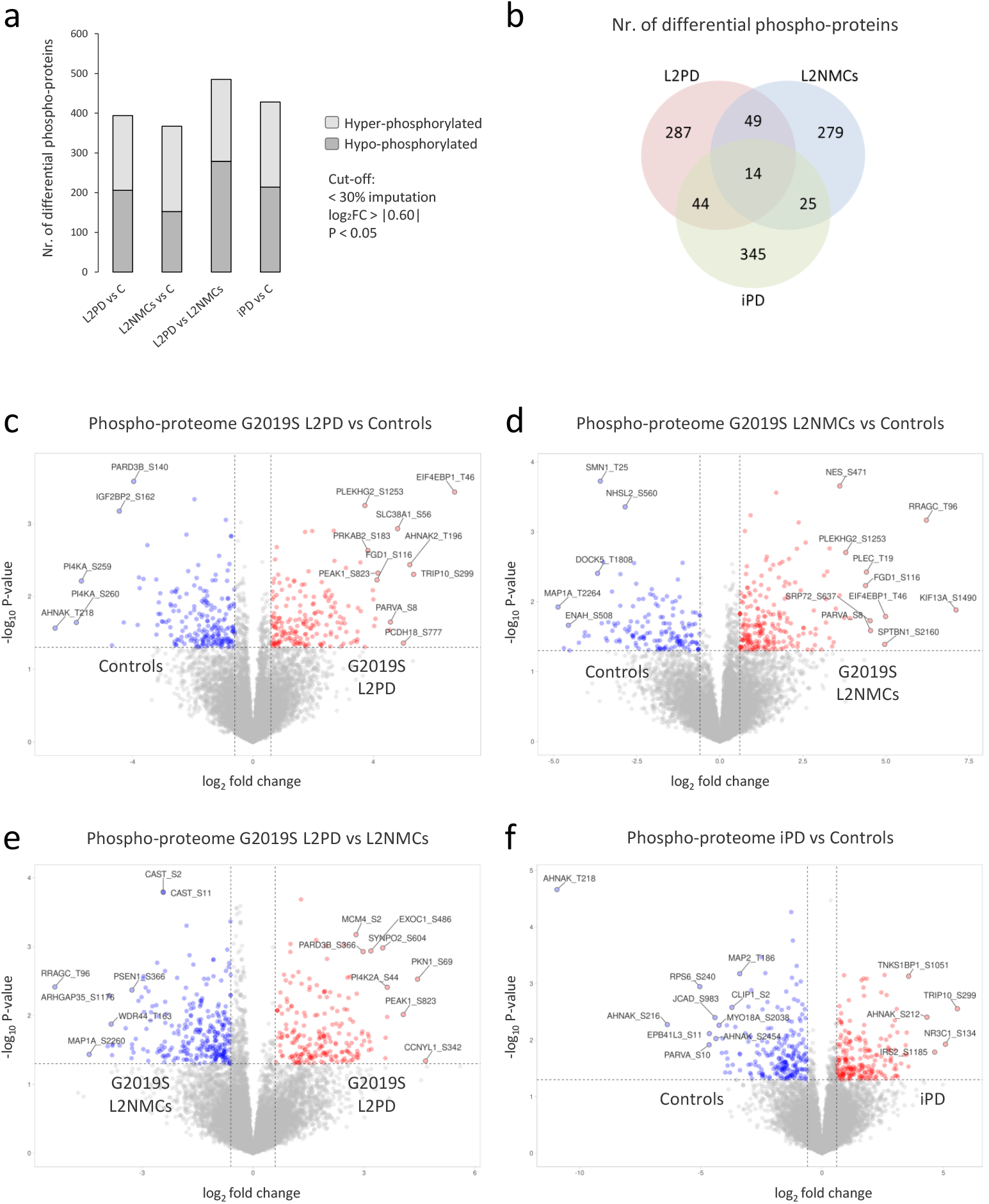
Phospho-proteome overview and differential analyses. **a)** Bar plot showing the number of differential phospho-proteins in different pairwise comparisons, with hyper-phosphorylated phospho-sites in dark grey and hypo-phosphorylated in light grey. The significance cut-off was set at log_2_FC≥⍰0.60⍰ and P<0.05. **b)** Venn diagram showing shared phospho-peptides between G2019S L2PD, G2019S L2NMCs, and iPD comparisons vs controls. **c)** Volcano plots showing phospho-site outcomes of the G2019S L2PD group compared to controls. Differential phospho-sites are shown in red (hyper-phosphorylated) and blue (hypo-phosphorylated). **d)** Phospho-site outcomes of the G2019S L2NMCs compared to controls. **e)** Phospho-site outcomes of the iPD group compared to controls. **f)** Phospho-site outcomes of the G2019S L2PD group compared to G2019S L2NMCs. Volcano plots were generated using default parameters in the VolcaNoSer software. Annotated dots correspond to the fifteen protein data points with the largest (Manhattan) distance from the origin above the significance thresholds indicated by the dashed line. VolcaNoSer: https://huygens.science.uva.nl/VolcaNoseR2/.

### LRRK2 phospho-proteome top differential hits

Among differential phospho-peptides, we identified elevated levels of pThr46 4E-BP1 as the most significant hit in G2019S L2PD vs controls (log_2_FC=6.7; P=3.7·10^−4^) and to a lesser extent in the L2NMC group as well (log_2_FC=3.2; P=9.2·10^−3^) (**Fig. 3**; **Fig. 4**). In addition, pThr37 4E-BP1 was also hyper-phosphorylated in iPD patients (log_2_FC=3.3; P-value 0.026). Importantly, two phosphorylation sites at 4E-BP1 (pThr37/46 4E-BP1) have been described as key regulators of the mTOR signalling pathway and essential for 40S ribosome cap-dependent translation, protein synthesis, and cell proliferation.^24,25^ Moreover, we identified another residue, pThr70 4E-BP1, which was also hyper-phosphorylated in L2PD patients (log_2_FC=1.8; P=0.04). The latter site represents a second checkpoint of 4E-BP1 activation dependent on prior pThr37/46 phosphorylation, which is required for 4E-BP1 release from eIF4E and initiation of mRNA translation into polypeptide chains.^25^ Other top phospho-proteins detected in L2PD included elevated levels of pSer823 PEAK1 and pSer1253 PLEKHG2^26^, which are related to Rho/Ras GTPase signalling and cytoskeleton regulation.^27,28^ In G2019S L2NMCs, we observed increased levels of pThr96 RRAGC, a key regulator of the cellular localisation of the mTOR complex.,^29^ and also of PLEKHG2 similar to L2PD. Lastly, iPD revealed deregulated hits such as elevated levels of pSer134 NR3C1, the glucocorticoid receptor, which is regulated by Akt/mTOR;^30^ hyper-phosphorylated levels of pSer1185 IRS2, which is related to insulin/Akt signalling; or down-regulated levels of pSer240 RPS6, a downstream protein target of the mTOR pathway.^31^ Overall, at the phospho-proteome level, we identified an mTOR pathway overactivation which affected known canonical phospho-proteins in G2019S carriers and also iPD (**Supplementary Fig. 1**).

**Figure 4:**
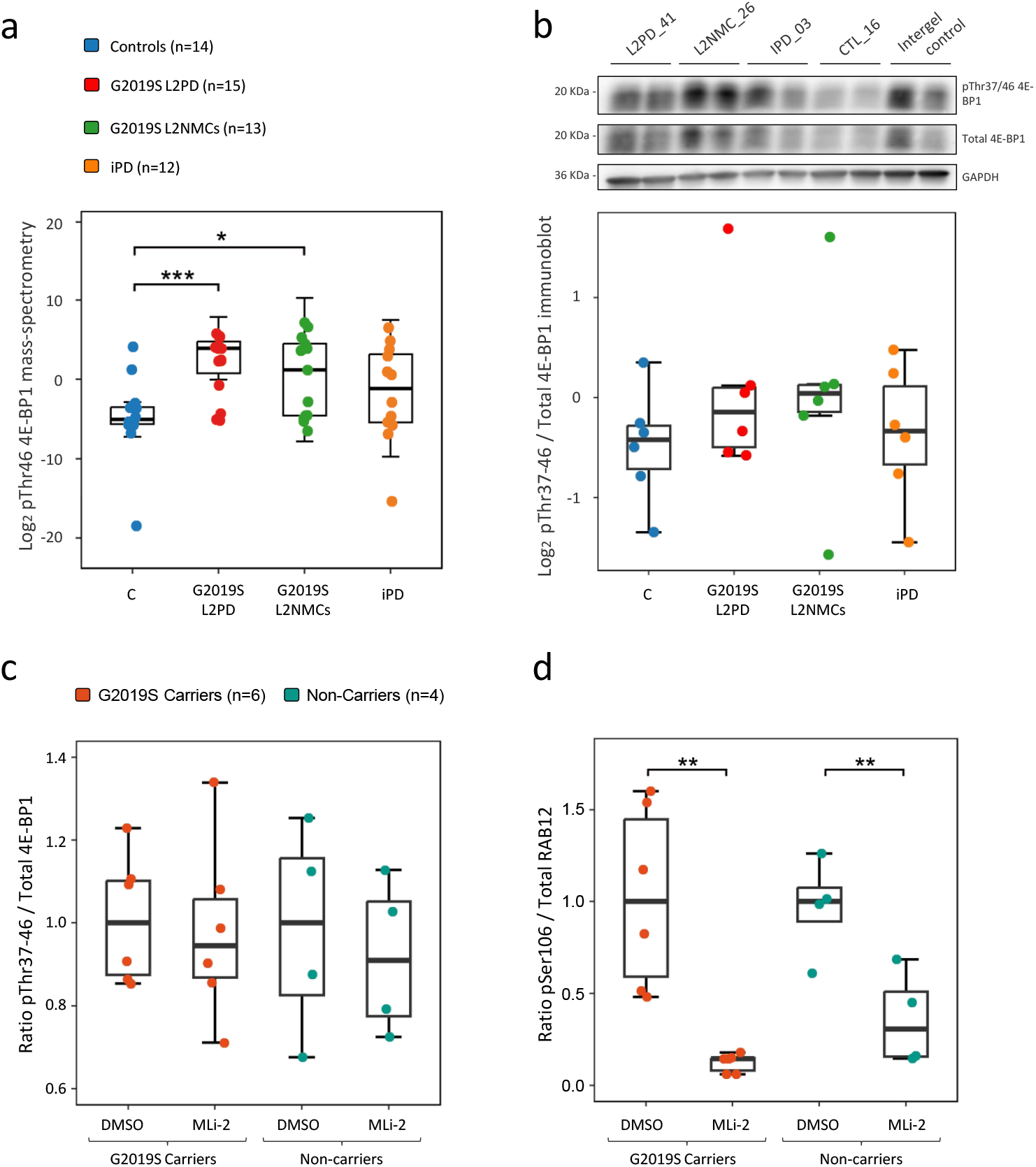
pThr46 4E-BP1 validation and MLi-2 inhibition assay. **a)** Boxplot representation of log_2_ pThr46 4E-BP1 phospho-proteome outcomes detected by DIA-MS of the entire cohort (n=54). **b)** Boxplot representation of log_2_ pThr37/46 / total 4E-BP1 phospho-protein levels obtained by immunoblot in a reduced cohort (n=24). Also, a representative blot is shown. Each sample was loaded twice, and each blot includes an intergel control to normalise between them. The entire analysis comprises six different blots, which are shown in the Supplementary data. **c)** Boxplot comparison of pThr37-46 / total 4E-BP1 phospho-protein levels between G2019S carriers and non-carriers treated with MLi-2 or not (n=10). **d)** Same comparison as previous, but with pSer106 / total RAB12 phosphoprotein levels. All blots are shown in the Supplementary data. * (p-value ≤ 0.05) ** (p-value ≤ 0.01) *** (p-value ≤ 0.001).

### Functional gene enrichment analysis of LRRK2 phospho-/proteomic differences

To identify the functional pathways and biological processes underlying the proteomic changes related to LRRK2 mutations, we stratified the differential hits from each comparison into up- and down-regulated proteins. We observed that down-regulated proteins in G2019S carriers compared to controls, either L2PD (65%) or L2NMCs (76%), were involved in mitochondrial energy metabolism and ribosomal protein synthesis (**Fig. 5**). Interestingly, we also observed very similar protein down-regulation (75%) effects in iPD, prominently affecting ribosomal proteins. In contrast, the differential proteome hits from G2019S L2PD compared to L2NMCs were mainly related to extracellular matrix/adhesion function (**Fig. 5**). On the other hand, up-regulated proteins in G2019S carriers vs controls, and also in iPD, were associated with endolysosomal function among other (**Supplementary Fig. 2**). At the phospho-proteome level, we applied the same approach and split the differential hits into hyper- and hypo-phospho-proteins. Of these, hyper-phosphorylated hits in G2019S carriers vs controls, and again also in iPD, were involved in Rho GTPase and mTOR signalling, with the strongest statistical significance observed for mTOR in G2019S L2PD and L2NMCs. Moreover, Rho GTPase, but not mTOR, was specifically enriched in G2019S L2PD vs L2NMCs (**Fig. 6**). Lastly, hypo-phosphorylated hits participated in Rho GTPase signalling in G20919S carriers and also in iPD, but not mTOR signalling, which was specific to hyper-phosphorylated hits (**Supplementary Fig. 3**). Overall, these results indicate that both LRRK2 G2019S carriers and iPD patients exhibit proteome changes mostly affecting mitochondrial and ribosomal functions, along with phospho-proteome deregulation involving the Rho GTPase and mTOR phospho-signalling pathways.

**Figure 5.**
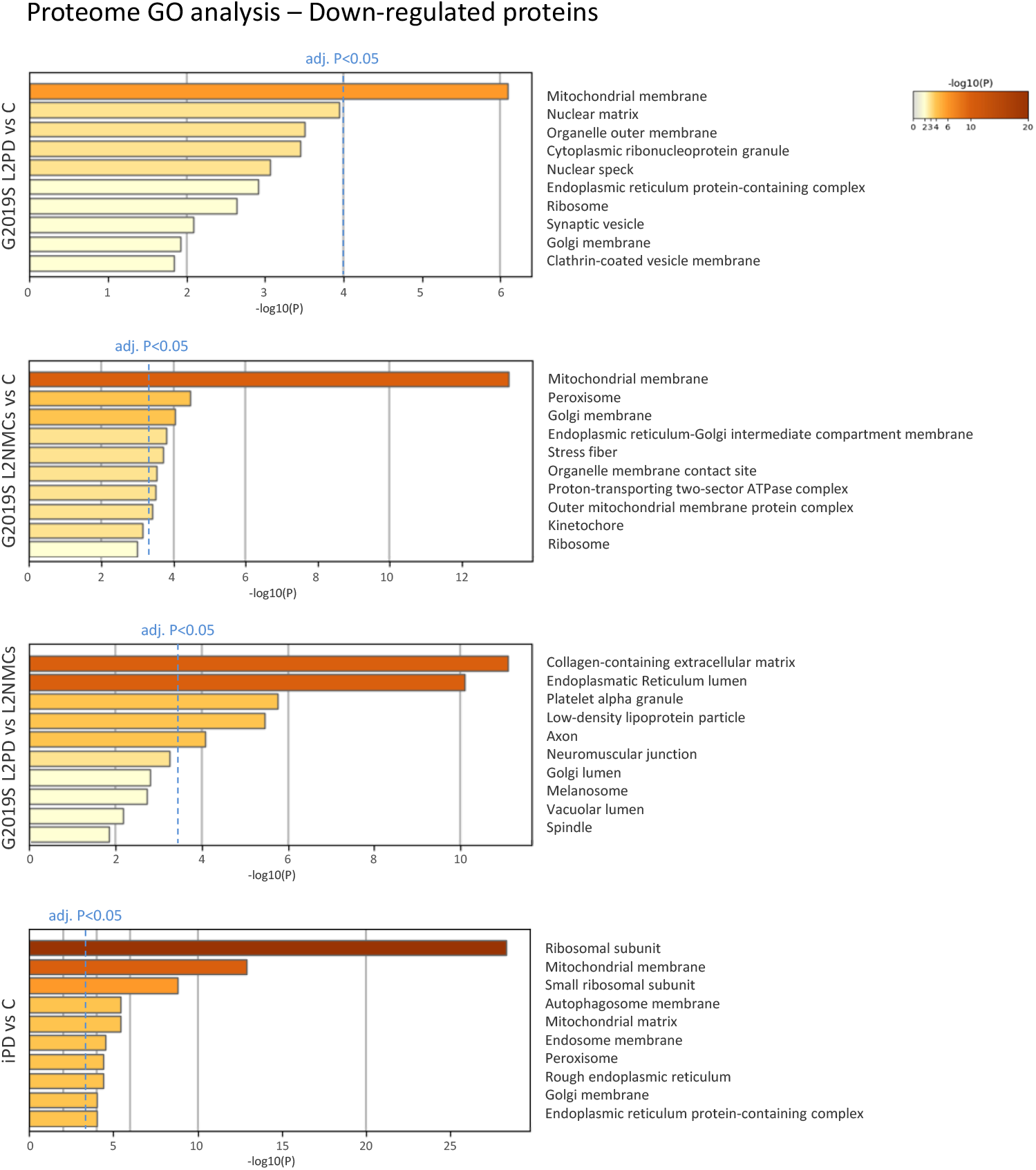
Biological enrichment analysis of the proteomic data. Biological enrichment analysis of down-regulated proteins in L2PD, L2NMCs and iPD compared with healthy controls, as well as L2PD compared with L2NMCs. Gene Ontology analysis for Cellular Components using Metascape with a *P*≤0.05. The blue-dashed line indicates the false discovery rate (FDR) adj. *P*≤0.05 cut-off.

**Figure 6.**
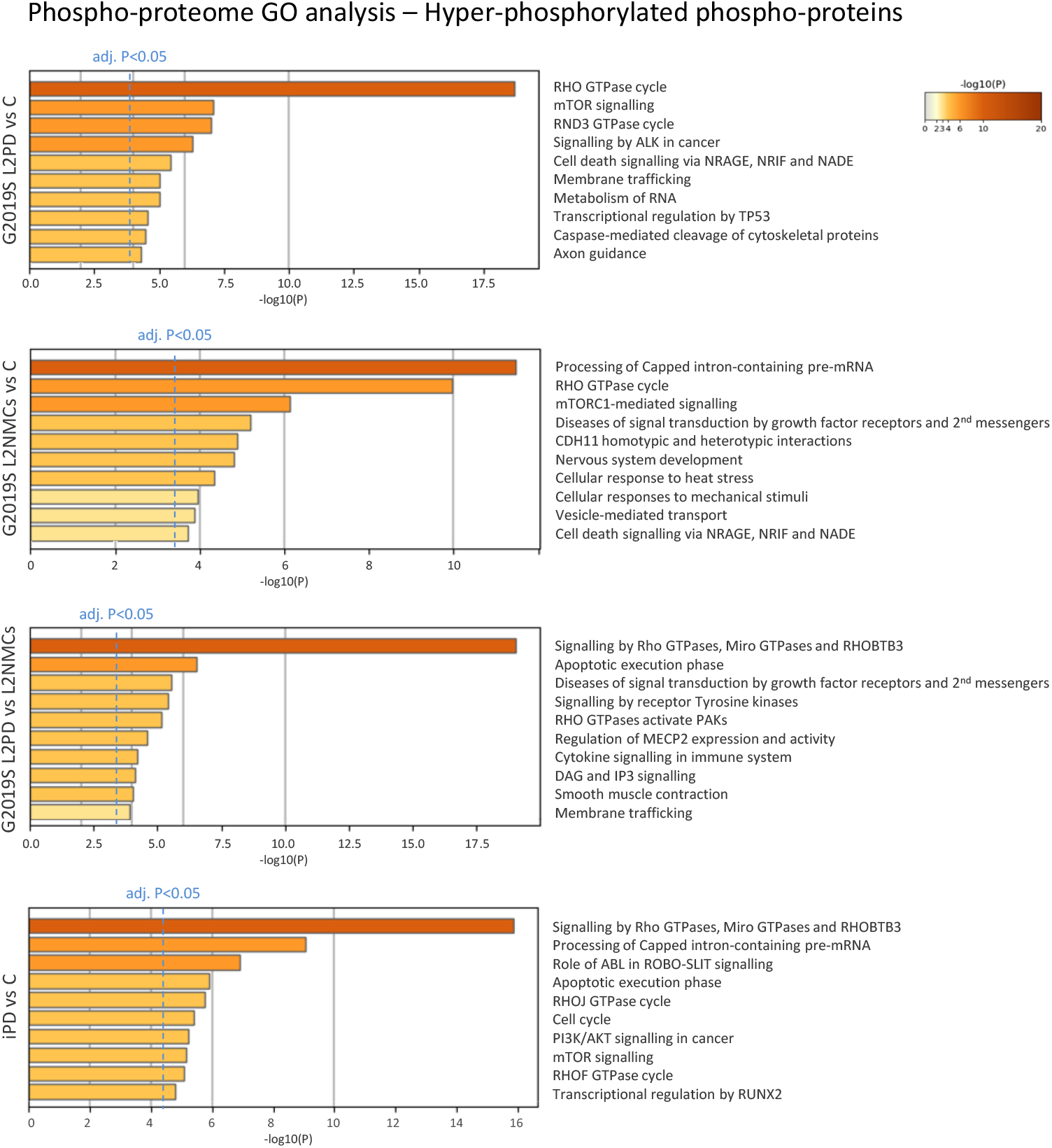
Biological enrichment analysis of the phospho-proteomic data. Biological enrichment analysis of hyper-phosphorylated phospho-proteins in L2PD, L2NMCs and iPD compared with controls, as well as L2PD compared with L2NMCs. Gene Ontology analysis for Reactome using Metascape with a cut-off of *P*≤0.05. The blue-dashed line indicates the false discovery rate (FDR) adj. *P*≤0.05 cut-off.

### Immunoblot assessment of endogenous phospho-/proteomic hits & LRRK2 inhibition

Using skin fibroblast cell lines from an additional set of participants (n=24), by immunoblot we assessed several the top-most differential hits detected in the DIA-MS analyses. Comparing DIA-MS and immunoblot data, we observed a pThr37/46 / total 4E-BP1 concordant expression pattern across groups, despite not reaching statistical significance (**Fig. 4**: **Supplementary Fig. 4**). However, other markers, such as ATG9 and MRPS14, did not show evident changes (**Supplementary Fig. 5**). This lack of statistical significance may be related to the lower sensitivity, dynamic range, and antibody limitations of antibody-based methods, such as immunoblot with regard gold-standard DIA-MS.^32^ In addition, we did not observe pThr37/46 4E-BP1 to total 4E-BP1 changes in response to MLi-2 LRRK2 inhibition, indicating that pThr37/46 4E-BP1 is not directly phosphorylated by LRRK2 under our experimental conditions (**Fig. 4**; **Supplementary Fig. 6**). As positive controls, we assessed the levels of pSer106 / total RAB12 (**Fig. 4**; **Supplementary Fig. 6**), pThr73 /total RAB10, and pSer935 / total LRRK2 (**Supplementary Fig. 7**). As expected, both pSer106 / total RAB12 and pThr73 / total RAB10 exhibited strong hypo-phosphorylation after MLi-2 treatment compared to non-treated samples (pSer106 RAB12 to total RAB12, paired T-test P=1.5×10^−4^; pThr73 RAB10 to total RAB10, paired T-test P=1.8×10^−4^), whereas pSer935 LRRK2 to total LRRK2 showed only subtle but significant down-regulation after LRRK2 inhibition (paired T-test P=0.014).

## DISCUSSION

To the best of our knowledge, this study represents, the first phospho-/proteome analysis of skin fibroblasts from a clinical LRRK2 cohort of PD manifesting and non-manifesting G2019S carriers.

At the proteome level, G2019S carriers exhibited significant protein deficits in both L2PD (65% down) and L2NMCs (76% down) as compared to controls, impacting mitochondrial energy and ribosomal protein synthesis and homeostasis. In these analyses, iPD shoed by far the largest amount of proteome differences, and similar to G2019S carriers, encompassed marked protein deficits (74% down), yet with a greater effect on ribosomal protein synthesis in iPD. Overall, protein down-regulation was earlier reported in LRRK2 peripheral blood mononuclear cells (PBMCs)^6,33^ and iPD fibroblasts.^12^ In addition, the observed mitochondrial protein deficits were in line with previous reports in PD models, underscoring the relevance of mitochondrial defects in PD.^34^ Altogether, these results indicate a pronounced mitochondria and ribosomal protein synthesis deficits in LRRK2 G2019S carriers, with and without PD symptoms, and also in iPD.

At the phospho-proteome level, hyper-phosphorylated proteins in both G2019S L2PD and L2NMCs primarily affected the mTOR and Rho GTPase signalling pathways, and interestingly, this was also the case in iPD. Whereas Rho GTPases regulate cytoskeletal dynamics, vesicle trafficking, and cell morphology, mTOR is a master regulator of protein synthesis and cell metabolism. Given that Rho GTPases can modulate mTOR signalling, these two signalling pathways have been shown to be functionally interconnected.^35^ Moreover, Rho GTPases^36,37^ and mTOR^38^ have been associated with LRRK2 activity, suggesting that their coordinated deregulation may contribute to pathogenic mechanisms in PD. Interestingly, the Rho GTPases pathway –but not mTOR– was significantly enriched in G2019S L2PD compared to G2019S L2NMCs. These findings suggest that mTOR deregulation represents an endogenous alteration occurring in PD, whereas Rho GTPase deregulation appears to evolve along with disease progression in G2019S carriers.

Here, it is important to note that defects at the phospho-proteome level of fibroblasts from our LRRK2 clinical cohort can be functionally related to changes at the proteome level. Specifically, it is well-established that mTOR can selectively regulate the translation of ribosomal and mitochondrial mRNAs in PD cell models, as well as in clinical studies.^39,40^ In addition, chronic LRRK2 inhibition using type-I small molecule LRRK2 inhibitors such as MLi-2 in the G2019S LRRK2 transgenic mice showed up-regulation of mitochondrial proteins compared to untreated controls,^41^ linking LRRK2 phospho-signalling to mitochondrial regulation and suggesting that LRRK2 may act upstream of mTOR. In line with this study, our findings in LRRK2 fibroblasts support that mTOR activation in G2019S LRRK2 carriers, symptomatic and asymptomatic, is associated with the alterations in mitochondrial and ribosomal proteins found at the proteome level, supporting a role of LRRK2 as one of the multiple regulators of the mTOR pathway.

mTOR pathway activation promotes protein translation, especially mitochondrial and ribosomal proteins.^24,42^ Consistent with these studies, enhanced mTOR signalling and protein synthesis have been reported in cellular and animal models of PD.^43^ However, earlier studies using primary PD fibroblasts have also reported mitochondrial, ribosomal and endolysosomal sorting protein down-regulation, similarly to our study.^12,44^ These results may reflect tissue-specific or temporal adaptive responses of the mTOR multi-function complex in controlling protein translation. For instance, chronic mTOR activation can lead to protein misfolding and endoplasmic reticulum stress, simultaneously triggering compensatory mechanisms, such as the mitochondrial integrated stress response (ISR)^45^ or other feedback loops^46^, ultimately leading to the deregulation of protein synthesis. Further studies investigating mTOR regulation in LRRK2 peripheral non-neuronal tissues beyond the CNS are needed to clarify this question.

RAB proteins are well-established substrates of mutant LRRK2,^47^ of which pSer106 RAB12^6^ and pThr73 RAB10^7^ have been validated in blood cells from LRRK2 clinical cohorts.^6^ In fibroblasts from G2019S carriers, we found elevated levels of pThr37 4E-BP1, which is a key downstream effector of mTOR.^24^ Previous studies have linked mutant LRRK2 activity to elevated pThr37/46 4E-BP1 phosphorylation and protein translation defects in *Drosophila* PD models;^38^ yet *in vitro* assays showed only weak 4E-BP1 phosphorylation effects by LRRK2.^48^ Moreover, elevated pThr37/46 4E-BP1 levels have also been reported in PD post-mortem brains.^43^ In our study, the overall concordance of pThr37/46 4E-BP1 findings in G2019S carriers by immunoblot and DIA-MS, along with the higher sensitivity of the latter, supports the biological validity of these findings. Yet, the lack of detectable LRRK2 inhibition on pThr37/46 4E-BP1 suggests that, unlike RAB proteins, indicates that 4E-BP1 is not a direct LRRK2 substrate or is only weakly phosphorylated by mutant LRRK2, as earlier described.^48^ Lastly, mTOR is regulated by multiple converging pathways, which may mask any potential individual effects of LRRK2 on pThr37/46 4E-BP1 levels.^49^

Despite the novel findings, our study has limitations. First, our analyses included only LRRK2 fibroblasts studied cross-sectionally at a single time point; however, future studies should investigate other peripheral cell types and tackle longitudinal experimental designs. Second, we have focused on the LRRK2 G2019S kinase mutations, but the phospho-proteomic effects of other LRRK2 pathogenic mutations, such as R1441G/C/H, warrant further investigation. Third, we did not assess potentially ongoing nigrostriatal degeneration in L2NMCs, for instance, by DaT-SPECT. Lastly, higher-resolution quantification methods and longer LRRK2 inhibition across different tissues may be necessary to detect subtle changes and clarify whether and how LRRK2 blockade modulates mTOR and Rho GTPases phospho-signalling in G2019S carriers.

Collectively, our findings have implications for future PD therapeutic strategies. Recently, a Mendelian randomisation study analysing the effect of mTOR effectors identified increased total 4E-BP1 levels in serum as protective for PD.^50^ However, despite the well-known neuroprotective effects of the mTOR inhibitor Sirolimus (rapamycin) in PD animal or cell models, its clinical translation remains limited.^51–53^ A recent phase Ib/IIa trial testing mTOR inhibitors in PD (RTB101 + sirolimus) was initiated but delayed due to the COVID-19 pandemic.^54^ Nevertheless, mTOR inhibitors have been shown to improve physiological parameters associated with ageing in both healthy individuals and those with ageing-related diseases.^55^ Taken together, these data underscore the role of 4E-BP1 in dopaminergic neuron protection and support further clinical exploration of modulators targeting the mTOR/4E-BP1 axis as a therapeutic approach for PD.

In summary, our study identifies deficits in ribosomal and mitochondrial proteins in LRRK2 G2019S carriers, which are also shared with iPD, and are related to aberrant mTOR signalling. These findings highlight the pivotal role of mTOR in regulating protein homeostasis at the peripheral level, with 4E-BP1 serving as a promising novel therapeutic target for future preclinical and clinical studies in PD. Further phospho-/proteomic analyses in additional LRRK2 clinical cohorts are needed to validate and expand these findings.

## METHODS

### Subjects

The study workflow is summarised in **Fig. 1**. The study was approved by the local ethics committee of the Hospital Clinic of Barcelona (code HCP/2020/0164), and conducted in accordance with the Declaration of Helsinki. Written informed consent was secured from all participants, and patient data was anonymised to protect confidentiality. Primary skin biopsies were collected from G2019S L2PD patients, G2019S L2NMCs, iPD patients, and control subjects. The cohort included n=15 G2019S L2PD (40% men, mean age at collection 64.7 ± 9.4 years (y.), mean age at onset 57.0 ± 10.4 y.), n=13 G2019S L2NMCs (38% men, mean age at collection 47.8 ± 9.6 y.), n=12 iPD (66% men, mean age at collection 58.6 ± 7.3 y., mean age at onset 51.3 ± 8.4 y.), and n=14 healthy controls (57% men, mean age at collection 62.9 ± 9.1 y.) (total n=54) (**Table 1**; **Supplementary Table 1**). We used Taqman assays (Thermo Fisher Sci. #C-63498123-10) with the Step One Plus Real-time PCR System (Life Tech. Inc.) to genotype LRRK2 G2019S and R1441G/C/H, as previously described.^3^ The L2PD and L2NMCs subjects were heterozygous carriers of G2019S, while iPD patients and healthy controls did not carry the LRRK2 G2019S or R1441G mutations. All study participants were of European ancestry.

### Fibroblast cell culture

3 mm diameter punch skin biopsies were taken from individuals from the alar surface of the arm. The biopsies were cut into approximately 1 mm pieces and cultured in 25 cm^2^ flasks using Dulbecco’s Modified Eagle’s Medium (DMEM) containing glucose (25⍰mM) (ThermoFisher Sci. #41966029), 10% heat-inactivated fetal bovine serum (FBS) (ThermoFisher Sci. #A5256801), and 1% Penicillin/Streptomycin (ThermoFisher Sci. #15140-122) at 37⍰°C and 5% CO_2_. The resulting dermal fibroblasts were expanded until confluence, trypsinised with 0,25% trypsin (ThermoFisher Sci. #15090046), and cryopreserved in 10% DMSO culture media in liquid N_2_ tanks until the experiment. Fibroblasts from all participants from the same passage number (**Supplementary Table 1**, mean cell passage of 1.4 ± 0.9) were cultured in parallel using the culture conditions described above until confluence, then harvested using trypsin/EDTA (ThermoFisher Sci. #15090-046) and centrifuged. Cell pellets were flash-frozen and stored at −80⍰°C until subsequent phospho-/protein isolation. The entire culture period for each subject ranged from 10 to 14 days, depending on the cell line.

### Sample preparation for DIA-MS

Fibroblast pellets from all study groups were processed concurrently. The experimental groups were blinded to the operator, and the runs were balanced according to randomised blind groups to reduce potential manipulation bias. Samples were homogenised in a lysis buffer comprising 7 M urea, 2 M thiourea, and 50 mM dithiothreitol (DTT), along with phosphatase and protease inhibitors. The lysates underwent centrifugation at 20.000 g for 1 hour at 15 °C, and the proteins in the supernatant were quantified using the Bradford assay kit (ThermoFisher Sci. #23200). Next, alkylation was performed by adding iodoacetamide to achieve a final concentration of 30 mM for 30 min in the dark at room temperature. The mixture was then diluted to 0.9 M urea using ammonium bicarbonate (ABC). After adding trypsin (Promega #TB309) at an enzyme-to-protein ratio of 1:20 (w/w), the samples were incubated at 37 °C for 18 hours. Digestion was halted by acidification (pH < 6) with acetic acid. Following the enzymatic cleavage of proteins, peptide cleaning was carried out using Pierce™ Peptide Desalting Spin Columns (ThermoFisher Sci. #89852). For total protein analyses, an amount equivalent to 10 µg of digested and cleaned peptides was separated after the desalting step and reconstituted to a final concentration of 200 ng/µL in 0.1% formic acid, with an internal reference transition (iRT) of 1:5000, before MS analysis. For phospho-peptide enrichment, 400⍰µg of protein were quantified and subjected to enzymatic digestion. Proteins were reduced by adding DTT to a final concentration of 20⍰mM and incubated at room temperature for 30 minutes. Phosphorylated peptides were enriched using the High-Select TiO_2_ Phospho-peptide Enrichment Kit (Thermo Fisher Sci. #A32993) according to the manufacturer’s protocol. Purification and concentration were carried out using C18 ZipTip Solid Phase Extraction (Millipore #ZTC18S). Phospho-peptides were then reconstituted to a final concentration of 37.5⍰ng/µL in 0.1% formic acid containing iRT peptides at a 1:20,000 dilution before MS analysis.

### DIA-MS analysis

Dried peptide samples were reconstituted in a solution of 2% acetonitrile (ACN) and 0.1% formic acid (FA) and quantified using a Nanodrop™ spectrophotometer (ThermoFisher Sci.), before Liquid Chromatography (LC) MS analysis. This analysis was conducted using an EASY-1000 nanoLC system connected to an Exploris 480 mass-spectrometer (Thermo Fisher Sci.). Peptides were separated through a C18 Aurora column (75 µm × 25 cm, 1.6 µm particles; IonOpticks) at a flow rate of 300 nL/min, utilising a 1 h gradient at 50 °c: 2% to 5% B in 1 min, 5% to 20% B in 48 min, 20% to 32% B in 12 min, and 32% to 95% B in 1 min (where A = 0.1% FA and B = 100% ACN:0.1% FA). Ionisation of peptides was achieved with a spray voltage of 1.6 kV at a capillary temperature of 275 °c. Sample data were collected in DIA mode, comprising full MS scans (scan range: 400 to 900 m/z; resolution: 60,000; maximum injection time: 22 ms; normalised AGC target: 3,000%) along with 24 periodic MS/MS segments, employing 20 Th isolation windows (0.5 Th overlap; resolution: 15,000; maximum injection time: 22 ms; normalised AGC target: 1,000%). Fragmentation of peptides was conducted using a normalised HCD collision energy of 30%.

### Phospho-/proteome differential analyses

Phospho-/proteome analyses refer to separate studies of total protein abundance (proteome) and protein phosphorylation of specific sites (phospho-proteome). The raw MS data files were analysed using Spectronaut (Biognosys) through direct data-independent analysis (dDIA). MS spectra were matched against the Uniprot human proteome reference database with standard settings. For proteome analysis, trypsin was used as the enzyme in a specific mode, with carbamidomethyl (C) designated as a fixed modification. In addition, oxidation (M), acetylation (protein N-term), deamidation (N), and Gln to pyro-Glu were defined as variable modifications for total protein analysis. In the phospho-proteome analysis, carbamidomethyl (C) remained a fixed modification, with oxidation (M), acetyl (protein N-term), and Phospho (STY) as variable modifications. Phospho-/protein identifications were filtered using a 1% Q-value. The resulting quantitative proteome data were exported to Perseus software (version 1.6.15.0) for statistical analysis and data visualisation. Missing values were imputed using the default settings in Perseus. We applied a width adjustment normalisation method and a log_2_ transformation on the contrast matrix for differential analysis, complemented by a limma statistical analysis. Only peptides with a P-value < 0.05 and a log_2_ fold change (FC) exceeding ⍰0.60⍰ were classified as differentially expressed. Some multiple test corrections in peptide results from proteome experiments may overlook actual positives, especially in studies with small sample sizes. Consequently, we employed dual thresholds, integrating statistical significance (*P*≤0.05) and biological relevance (|log_2_FC|≥0.60), rather than applying stringent multiple-testing corrections (e.g., the Bonferroni or false discovery rate [FDR] method). This approach was chosen given the moderate sample size and exploratory nature of the study, as overly conservative corrections can obscure biologically meaningful differences in proteomic data.^56^ Quantitative phospho-/proteome data were integrated using a custom-coded Peptide Collapse plugin (v. 1.1.4.4) within Perseus (1.6.15.0) to convert a standard Spectronaut report into a site-level report. The default plugin settings grouped post-translational modifications (PTMs) by sample, collapsed the matrix at the site level, and applied a PTM localisation probabilities filter with a threshold of greater than 0.75. Statistical analyses were performed following the same protocol as in the proteome study.

### Functional enrichment of LRRK2 fibroblasts

Enrichment analyses of the differentially phosphorylated proteins detected across various group comparisons were conducted using Gene Ontology (GO) analysis in Metascape.^57^ We used a less stringent cut-off (P≤0.05; log_2_FC ≥⍰0.38⍰) to ensure having enough data to perform gene enrichment and increase the range of GO terms, thereby ensuring sufficient power for enrichment analysis. We used the Metascape settings, including a minimal gene overlap of 3, a minimum fold enrichment of |1.5|, and a P-value cut-off of P<0.05. Results were sorted by Benjamini & Hochberg false discovery rate (FDR) multiple-testing adjusted P<0.05. Specifically, Cellular Component GO terms were used for the proteome analysis, stratified by up- and down-regulated proteins, while Reactome pathway annotations were applied to the phospho-proteome, stratified by hyper- and hypo-phosphorylated phospho-peptides.

### Immunoblot assessment of LRRK2 differential phospho-/proteins

We newly cultured six fibroblast cell lines from each group (G2019S L2PD, G2019S L2NMCs, iPD, and controls; see **Supplementary Table 2**). Fibroblasts were seeded at matching cell passage numbers all below pass number 7 (mean cell passage of 4.4 ± 1.1), grown until confluence, and passed using trypsin/EDTA (Sigma #15090-046). After reaching the second confluence, the cells were harvested, and dry pellets were stored at −80⍰°C until subsequent protein extraction. Cold lysis buffer was added to each pellet, and the mixture was incubated on ice for 10 min, with regular vortexing of the microcentrifuge tube. Samples were centrifuged at 14,000 x g for 15 min to collect the cell debris, and the supernatant was transferred to a new tube for quantification analysis. Sample protein concentrations were determined using the Pierce™ Coomassie (Bradford) (ThermoFisher Sci. #23200). Cell extracts were mixed with a quarter of a volume of 4x Laemmli Sample Buffer (BioRad #1610747) and 1% of 2-mercaptoethanol. Samples were heated at 95°C for 5 min. 30 μg of each sample was loaded onto 4–20% Mini-PROTEAN® TGX™ Precast Protein Gels (BioRad #4561094) and electrophoresed with 10x Tris/Glycine/SDS Electrophoresis Buffer (BioRad #1610772EDU). After electrophoresis, proteins were transferred onto a 0.45 µm nitrocellulose membrane (BioRad #1620115) at 100 V for 1 h. The transferred membrane was blocked with 5% (w/v) bovine serum albumin dissolved in TBS-T [20 mM Tris-HCl, pH 7.5, 150 mM NaCl, and 0.1% (v/v) Tween 20] at room temperature for 1 h. Each sample was loaded in duplicate, and the different antibodies (summarised in **Supplementary Table 3**) were arranged in two identical membranes, one for 4E-BP1/GAPDH and the other for ATG9a/MRPS14/GAPDH. Membranes were incubated with the primary antibody in 5% (w/v) bovine serum albumin in TBS-T overnight at 4°C. Before the secondary antibody incubation, membranes were washed three times with TBS-T for 5 min each. The secondary antibody was prepared at a 1:10,000 dilution in 5% (w/v) bovine serum albumin in TBS-T and incubated for 1 h at room temperature. Membranes were washed with TBS-T three times, with a 10-minute incubation for each wash. Protein bands were acquired using the LAS4000 imaging system and quantified using the Image Studio software, and significance was determined by the Wilcoxon signed-rank test (*P* ≤ 0.05).

### MLi-2 LRRK2 inhibition

We cultured another subset of fibroblast lines from n=10 subjects to test the effect of the LRKK2 kinase inhibitor MLi-2 to measure potential phosphorylation changes in the candidate target pThr37/46 4E-BP1. The subset encompassed fibroblasts from G2019S L2PD (n=4), G2019S L2NMCs (n=2), iPD (n=2) and healthy controls (n=2), seeded at matching cell passage numbers all below pass number 6 (mean cell passage of 4.5 ± 1.1) (**Supplementary Table 4**). Each fibroblast line was harvested and seeded onto two 100 mm dishes (ThermoFisher Sci. #150464), with 300,000 cells per dish, to perform⍰LRRK2 pharmacological inhibition with MLi-2, using technical replicates. Briefly, each technical replicate was treated with either 100 nM MLi-2 or an equivalent volume of DMSO for 1h at room temperature. After the treatment, the culture medium containing MLi-2 or DMSO was aspirated, and cells were washed with PBS. Cell lysis buffer (Cell Signalling Tech. #cst9803S) contained 20 mM Tris-HCl (pH 7.5), 150 mM NaCl, 1 mM Na_2_EDTA, 1 mM EGTA, 1% Triton, 2.5 mM sodium pyrophosphate, 1 mM beta-glycerophosphate, 1 mM Na_3_VO_4_ and 1 µg/ml leupeptin, and was supplemented with 1 mM phenyl-methyl-sulfonyl fluoride (PMSF) (Sigma #10837091001), cOmplete protease (Roche #11697498001) and PhosSTOP phosphatase (Roche #4906845001) inhibitors. This cell lysis buffer was added to the plates and incubated on ice for 10 min. After incubation, cells were scraped, collected into microcentrifuge tubes, and centrifuged at 14,000 x g at 4°C for 10 min. The supernatants were stored at −80°C until used. The immunoblotting procedure was the same as that described above. Technical replicates for the MLi-2 treatment were included by loading duplicate samples onto the gel. Samples were also arranged in two identical membranes, one for RAB12/4E-BP1/GAPDH antibodies and the other for LRRK2/RAB10/GAPDH (**Supplementary Table 3**). For RAB12/4E-BP1/GAPDH membranes, the blots were performed in triplicate.

## ABBREVIATIONS

(Minimal abbreviations to be used in the text)

DIA: Data independent acquisition
MS: Mass-spectrometry
PD: Parkinson’s disease
LRRK2: Leucine-rich repeat kinase 2
L2PD: G2019S LRRK2-associated PD patients
L2NMCs: G2019S LRRK2 disease non-manifesting carriers
iPD: Idiopathic PD patients
p: Phospho-
FC: Fold-change
CNS: Central nervous system

## DATA AVAILABILITY

The mass spectrometry proteomics data have been deposited in the PRIDE partner repository with the dataset identifiers PXD055255 for the proteome and PXD055342 for the phospho-proteome analyses. In addition, to promote transparency and open science, phospho-/proteomic results from this study are openly available through the Curtain software platform. This user-friendly, web-based tool enables interactive exploration of significant hits with different cut-offs, the generation and visualisation of volcano plots, and non-imputed bar plots of individual hits. The analysis can be accessed and reproduced via the following link(s): proteome comparisons <u>G2019S L2PD vs C</u>, <u>G2019S L2NMCs vs C</u>, <u>G2019S L2PD vs L2NMCs</u>, and <u>iPD vs C</u>; phospho-proteome comparisons <u>G2019S L2PD vs C</u>, <u>G2019S L2NMCs vs C</u>, <u>G2019S L2PD vs L2NMCs</u>, and <u>iPD vs C</u>.

## ACKNOWLEDGEMENTS

We thank the patients and their relatives for their continued and essential collaboration. We are grateful to Toan K. Phung for his assistance in preparing and adapting the dataset for integration into the Curtain platform.

## FUNDING

This work was supported by the “Proyectos I+D+I en Salud” programme from the Instituto de Salud Carlos III (ISCIII) (#PIE20/00259) to M.E., co-funded by the European Union. R.F.S. was supported by a Miguel Servet grant (#CP19/00048) and two “Proyectos I+D+I en Salud” grants (#PI20/00659 and #PI23/00661) from the ISCIII, co-funded by the European Union. A.R.C. was funded by the EU Next-Generation 2022 Investigo programme from the European Commission (EC) / Agència de Gestió d’Ajuts Universitaris i de Recerca (AGAUR) (#100028TC2) and the “Proyectos I+D+I en Salud” programme (#PI23/00661) from the ISCIII, co-funded by the European Union, awarded to R.F.S. IDIBAPS receives support from the CERCA Programme of the Generalitat de Catalunya. M.F. was funded by the María de Maeztu Programme (MICIU/AEI #CEX2021-001159-M) awarded to the Parkinson’s Disease and Movement Disorders Group of the Institut de Neurociències, Universitat de Barcelona.

## AUTHOR CONTRIBUTIONS

Funding acquisition: M.E. Study Conception: M.E., R.F.S. Work supervision: M.E., R.F.S. Phospho-/proteomic analyses: J.F.I., A.C., E.S. Western blot experiments: A.R.C., G.P. Skin biopsy recruitment: M.JM., A.G., M.F., F.V., C.P., A.S.G., Y.C., E.T., A.C. Fibroblast culture: A.R.C., M.F., G.P. Statistical Analysis: J.F.I., A.R.C. Writing of the first draft: A.R.C., M.E. Figure and tables preparation: A.R.C., M.E., R.F.S. Review and Critique: all authors. All authors read and approved the final manuscript.

## TABLE & FIGURE LEGENDS

**Table 1. Participant clinic-demographics.** Data expressed as a mean ± standard deviation (S.D.) with the number of available subjects/totals in brackets. L2PD = LRRK2-associated PD patients; L2NMCs = LRRK2 non-manifesting carriers; iPD = idiopathic PD; C = controls; AAO = age-at-onset; “-” = not applicable.

## Notes

### Competing Interest Statement

The authors have declared no competing interest.

