## Supplementary material for "Phospho-signalling deregulation of the mTOR pathway in fibroblasts from LRRK2 Parkinson’s patients": Supplementary tables and figures.pdf

**Supplementary Table 1: Summary of subject clinical features from Parkinson disease (PD) patients, carriers and controls.** L2PD = LRRK2-associated PD patients; L2NMCs = LRRK2 non-manifesting carriers; iPD = idiopathic PD; C = controls; AAO = age-at-onset.

| Patient Code | Code | Group | Familial PD | Sex | AAD | AAO | Cell passage | DAT scan+ |
|---|---|---|---|---|---|---|---|---|
| FOX-002-003 | L2PD_38 | L2PD | Yes | female | 75 | 71 | px1 | NA |
| FOX-003-001 | L2PD_39 | L2PD | Yes | male | 56 | 40 | px1 | NA |
| FOX-004-002 | L2PD_40 | L2PD | Yes | male | 66 | 64 | px1 | NA |
| FOX-019-001 | L2PD_09 | L2PD | Yes | male | 71 | 58 | px1 | NA |
| FOX-038-001 | L2PD_37 | L2PD | Yes | male | 70 | 70 | px1 | NA |
| FOX-039-001 | L2PD_36 | L2PD | Yes | female | 77 | 66 | px1 | NA |
| FOX-045-001 | L2PD_35 | L2PD | Yes | male | 77 | 67 | px1 | NA |
| FOX-049-001 | L2PD_34 | L2PD | Yes | female | 59 | 57 | px1 | NA |
| FOX-050-001 | L2PD_33 | L2PD | not known | female | 64 | 58 | px1 | NA |
| FOX-068-001 | L2PD_13 | L2PD | Yes | female | 58 | 58 | px1 | NA |
| FOX-072-001 | L2PD_41 | L2PD | Yes | female | 72 | 60 | px1 | NA |
| FOX-075-001 | L2PD_08 | L2PD | Yes | male | 51 | 46 | px1 | NA |
| SP-06 | L2PD_10 | L2PD | Yes | male | 44 | 34 | px3 | NA |
| SP-12 | L2PD_11 | L2PD | Yes | female | 63 | 49 | px2 | NA |
| SP-13 | L2PD_12 | L2PD | Yes | female | 68 | 57 | px4 | NA |
| BIOP-202 | L2NMC_54 | L2NMCs | Yes | female | 63 | NA | px1 | Yes |
| FOX-004-008 | L2NMC_21 | L2NMCs | Yes | male | 40 | NA | px1 | Yes |
| FOX-004-010 | L2NMC_22 | L2NMCs | Yes | male | 37 | NA | px1 | No |
| FOX-005-001 | L2NMC_23 | L2NMCs | Yes | male | 66 | NA | px1 | Yes |
| FOX-023-003 | L2NMC_24 | L2NMCs | Yes | female | 61 | NA | px1 | Yes |
| FOX-023-007 | L2NMC_25 | L2NMCs | Yes | female | 47 | NA | px1 | No |
| FOX-023-008 | L2NMC_26 | L2NMCs | Yes | male | 50 | NA | px1 | No |
| FOX-023-009 | L2NMC_27 | L2NMCs | Yes | female | 43 | NA | px1 | No |
| FOX-039-002 | L2NMC_49 | L2NMCs | Yes | female | 51 | NA | px1 | No |
| FOX-068-002 | L2NMC_52 | L2NMCs | Yes | male | 34 | NA | px1 | No |
| FOX-071-002 | L2NMC_51 | L2NMCs | Yes | female | 44 | NA | px1 | No |
| FOX-075-002 | L2NMC_50 | L2NMCs | Yes | female | 45 | NA | px1 | No |
| MF-006-01 | L2NMC_53 | L2NMCs | Yes | female | 41 | NA | px1 | NA |
| FOX-015-001 | IPD_01 | iPD | No | male | 68 | 56 | px1 | NA |
| IPD-008 | IPD_03 | iPD | No | female | 65 | 57 | px1 | NA |
| IPD-009 | IPD_28 | iPD | not known | male | 66 | 61 | px1 | NA |
| IPD-010 | IPD_29 | iPD | No | male | 62 | 53 | px1 | NA |
| IPD-013 | IPD_30 | iPD | No | female | 47 | 41 | px1 | NA |
| IPD-014 | IPD_31 | iPD | not known | male | 59 | 58 | px1 | NA |
| MF-012-01 | IPD_32 | iPD | No | male | 58 | 35 | px1 | NA |
| SP-01 | IPD_04 | iPD | No | female | 62 | 59 | px3 | NA |
| SP-02 | IPD_05 | iPD | No | male | 54 | 48 | px3 | NA |
| SP-04 | IPD_06 | iPD | No | male | 46 | 40 | px2 | NA |
| SP-08 | IPD_02 | iPD | not known | female | 66 | 60 | px1 | NA |
| SP-16 | IPD_07 | iPD | sporadic | female | 51 | 48 | px2 | NA |
| CTRL-001 | CTL_16 | Control | No | male | 68 | NA | px1 | NA |
| CTRL-002 | CTL_18 | Control | No | female | 73 | NA | px1 | NA |
| CTRL-003 | CTL_15 | Control | No | female | 55 | NA | px1 | NA |
| CTRL-004 | CTL_44 | Control | No | female | 55 | NA | px1 | NA |

| | | | | | | | | |
|---|---|---|---|---|---|---|---|---|
| FOX-002-001 | CTL_48 | Control | No | male | 53 | NA | px1 | NA |
| FOX-018-002 | CTL_46 | Control | No | male | 83 | NA | px1 | NA |
| FOX-024-002 | CTL_47 | Control | No | male | 67 | NA | px1 | NA |
| FOX-050-002 | CTL_45 | Control | No | male | 69 | NA | px1 | NA |
| FOX-069-001 | CTL_14 | Control | No | male | 64 | NA | px1 | NA |
| FOX-073-001 | CTL_43 | Control | No | female | 66 | NA | px1 | NA |
| FOX-076-001 | CTL_17 | Control | No | female | 61 | NA | px1 | NA |
| SP-09 | CTL_19 | Control | No | male | 66 | NA | px4 | NA |
| SP-11 | CTL_20 | Control | No | female | 48 | NA | px3 | NA |
| SP-17 | CTL_42 | Control | No | male | 52 | NA | px5 | NA |

**Supplementary Table 2. Summary of subject clinical features from Parkinson disease (PD) patients, carriers and controls used for the immunoblot assessment.** L2PD = LRRK2-associated PD patients; L2NMCs = LRRK2 non-manifesting carriers; iPD = idiopathic PD; C = controls; AAO = age-at-onset.

| Patient Code | Code | Group | Sex | Cell passage |
|---|---|---|---|---|
| FOX-004-002 | L2PD_40 | L2PD | male | px3 |
| FOX-019-001 | L2PD_09 | L2PD | male | px5 |
| FOX-038-001 | L2PD_37 | L2PD | male | px5 |
| FOX-045-001 | L2PD_35 | L2PD | male | px3 |
| FOX-050-001 | L2PD_33 | L2PD | female | px4 |
| FOX-072-001 | L2PD_41 | L2PD | female | px6 |
| FOX-004-008 | L2NMC_21 | L2NMCs | male | px3 |
| FOX-023-003 | L2NMC_24 | L2NMCs | female | px3 |
| FOX-023-007 | L2NMC_25 | L2NMCs | female | px4 |
| FOX-023-008 | L2NMC_26 | L2NMCs | male | px7 |
| FOX-039-002 | L2NMC_49 | L2NMCs | female | px6 |
| FOX-071-002 | L2NMC_51 | L2NMCs | female | px3 |
| IPD-008 | IPD_03 | iPD | female | px4 |
| IPD-009 | IPD_28 | iPD | male | px4 |
| IPD-010 | IPD_29 | iPD | male | px4 |
| IPD-013 | IPD_30 | iPD | female | px4 |
| IPD-014 | IPD_31 | iPD | male | px4 |
| SP-01 | IPD_04 | iPD | female | px5 |
| CTRL-001 | CTL_16 | Control | male | px5 |
| CTRL-002 | CTL_18 | Control | female | px4 |
| FOX-050-002 | CTL_45 | Control | male | px5 |
| FOX-069-001 | CTL_14 | Control | male | px4 |
| FOX-073-001 | CTL_43 | Control | female | px5 |
| FOX-076-001 | CTL_17 | Control | female | px6 |

**Supplementary Table 3: Summary of antibodies used for immunoblotting**

| Antibody | Brand | Reference | Host specie | Dilution |
|---|---|---|---|---|
| pSer935 LRRK2 | Abcam | ab133450 | Rabbit | 1:500 |
| LRRK2 | Antibodies | A305041 | Mouse | 1:250 |
| pThr37/46 4EBP1 | Cell Signalling | 2855 | Rabbit | 1:1000 |
| 4EBP1 | ThermoFisher | AHO1382 | Mouse | 1:250 |
| ATG9a | Abcam | ab108338 | Rabbit | 1:500 |
| MRPS14 | Protein Tech | 16301-1-AP | Rabbit | 1:1000 |
| pThr73 Rab10 | Abcam | MJF-R21 | Rabbit | 1:1000 |
| Rab10 | NanoTools | 0680-100 | Mouse | 1:1000 |
| pSer106 Rab12 | Abcam | ab256487 | Rabbit | 1:500 |
| Rab12 | Santa Cruz | sc-515613 | Mouse | 1:500 |
| GAPDH | Santa Cruz | Sc-32233 | Mouse | 1:4000 |
| Anti-Mouse IgG H&L (HRP) | Abcam | ab205719 | Goat | 1:10000 |
| Anti-Rabbit IgG H+L (HRP) | Invitrogen | 31460 | Goat | 1:10000 |

**Supplementary Table 4. Summary of subject clinical features from Parkinson disease (PD) patients, carriers and controls used for the inhibition study.** L2PD = LRRK2-associated PD patients; L2NMCs = LRRK2 non-manifesting carriers; iPD = idiopathic PD; C = controls; AAO = age-at-onset.

| Patient Code | Code | Group | Sex | Cell passage |
|---|---|---|---|---|
| FOX-045-001 | L2PD_35 | L2PD | male | px4 |
| FOX-050-001 | L2PD_33 | L2PD | female | px5 |
| FOX-019-001 | L2PD_09 | L2PD | male | px5 |
| FOX-072-001 | L2PD_41 | L2PD | female | px6 |
| FOX-004-008 | L2NMC_21 | L2NMCs | male | px5 |
| FOX-071-002 | L2NMC_51 | L2NMCs | female | px2 |
| IPD-010 | IPD_29 | iPD | male | px4 |
| IPD-014 | IPD_31 | iPD | male | px5 |
| CTRL-003 | CTL_15 | Control | female | px4 |
| FOX-073-001 | CTL_43 | Control | female | px5 |

**Supplementary Figure 1. Phospho-peptides affecting the mTOR pathway in L2PD, L2NMCs, or iPD groups compared to healthy controls.** Down-stream phospho-proteins of mTORC are marked in green and up-stream in blue. An arrow indicates activation according to the bibliography. The dashed line indicates an indirect effect.

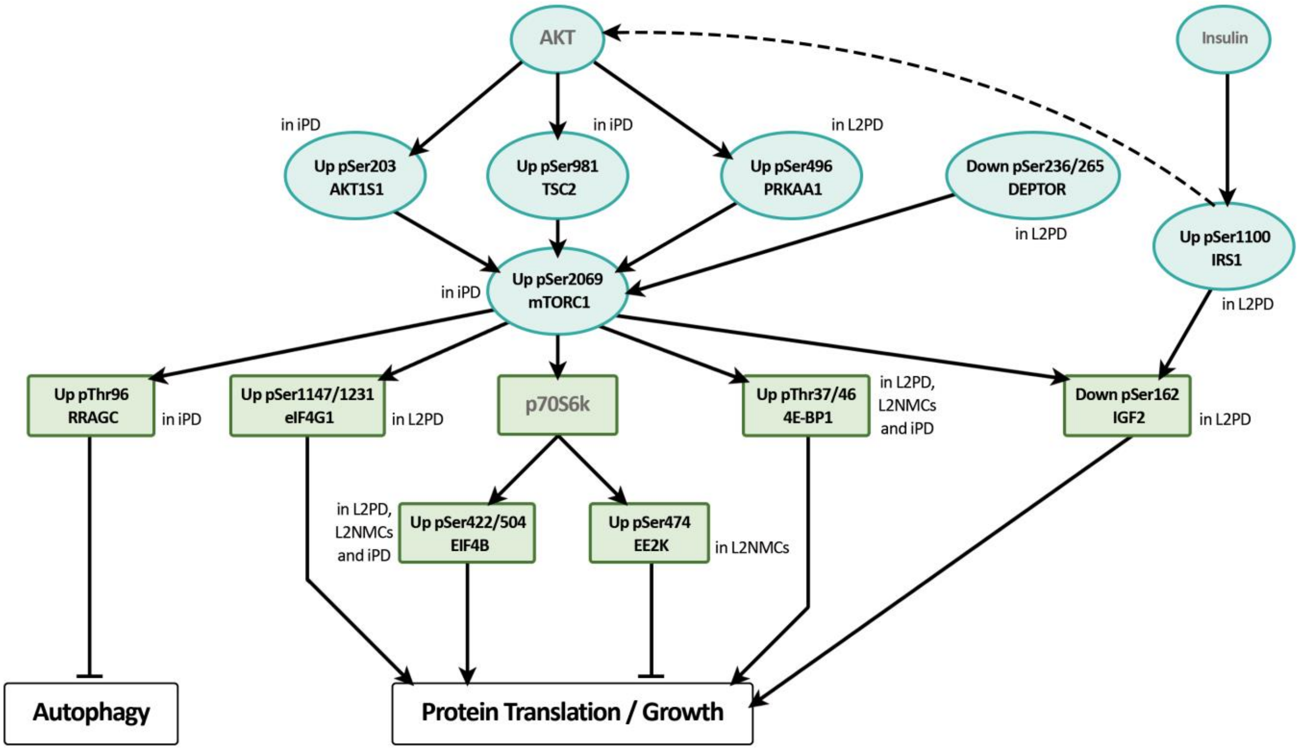


### Bibliography used to build phosphorylation signalling representation


- pSer981 TSC2 (PMID: 16636147)
- pSer496 PRKAA1 (PMID: 32130880)
- pSer203 AKT1S1 (PMID: 26876154)
- pSer236/265 DEPTOR (PMID: 22745583)
- pSer1147/1231 eIF4G1 (PMID: 35163752)
- pSer422/504 EIF4B (PMID: 23105104)
- pThr37/46 4E-BP1 (PMID: 10364159)
- pSer112 4E-BP1 (PMID: 11146653)
- pSer474 EF2K (PMID: 32859902)
- pThr96 RRAGC (PMID: 30552228)
- pSer162 IGF2BP2 (PMID: 21576258)
- pSer1100 IRS1 (PMID: 32444657)

**Supplementary Figure 2. Biological enrichment analysis of the proteomic data.** Biological enrichment analysis of up-regulated proteins in L2PD, L2NMCs and iPD compared with healthy controls, as well as L2PD compared with L2NMCs. Gene Ontology analysis for Cellular Components using Metascape with a cut-off of P-value ≤ 0.05. The blue dashed line indicates the false discovery rate (FDR) adj. P ≤ 0.05 cut-off.

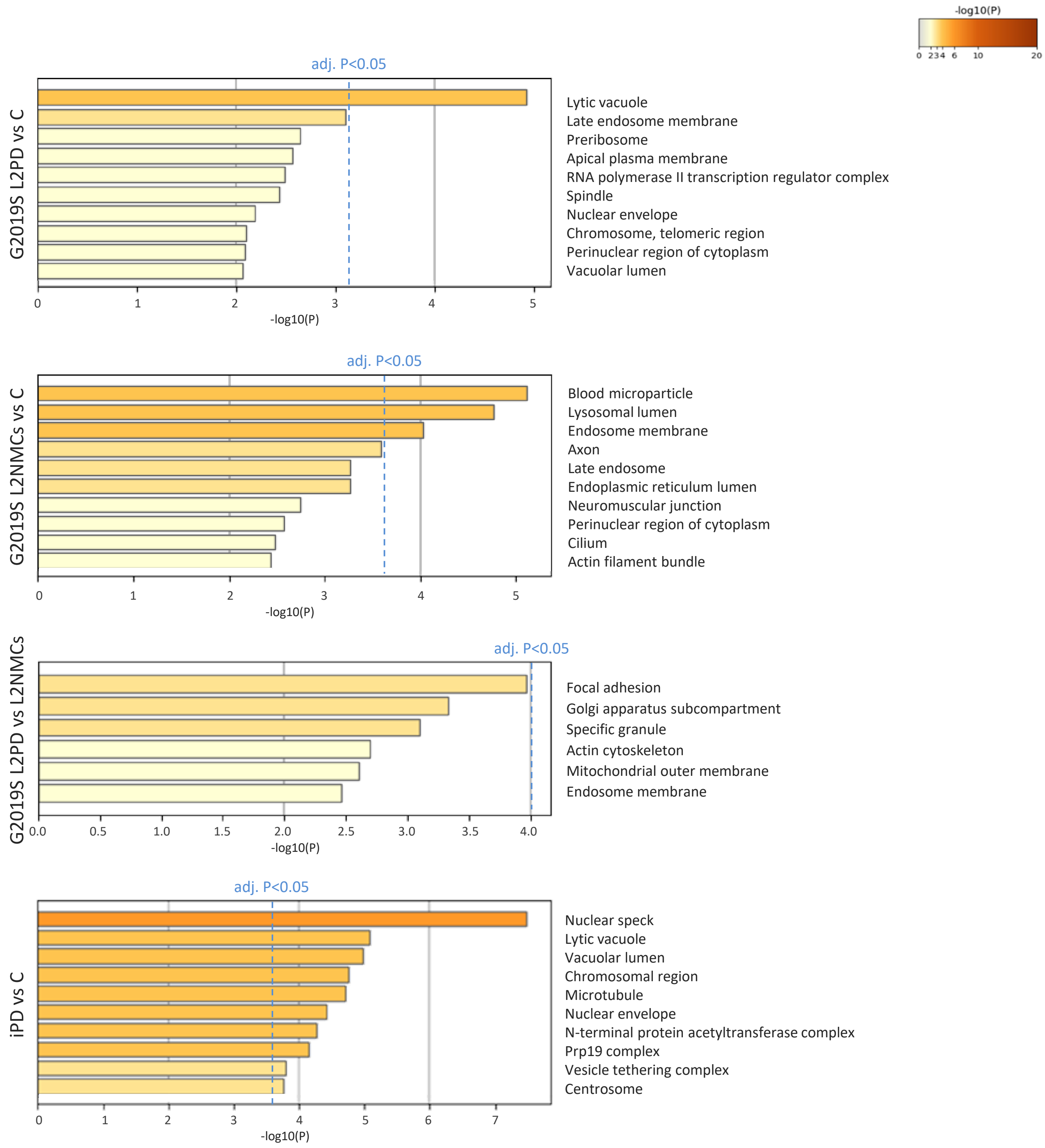

**Supplementary Figure 3. Biological enrichment analysis of the phospho-proteomic data.** Biological enrichment analysis of hypo-phosphorylated phospho-proteins in L2PD, L2NMCs and iPD compared with controls, as well as L2PD compared with L2NMCs. Gene Ontology analysis for Reactome using Metascape with a cut-off of P-value ≤ 0.05. The blue dashed line indicates the false discovery rate (FDR) adj. P ≤ 0.05 cut-off.

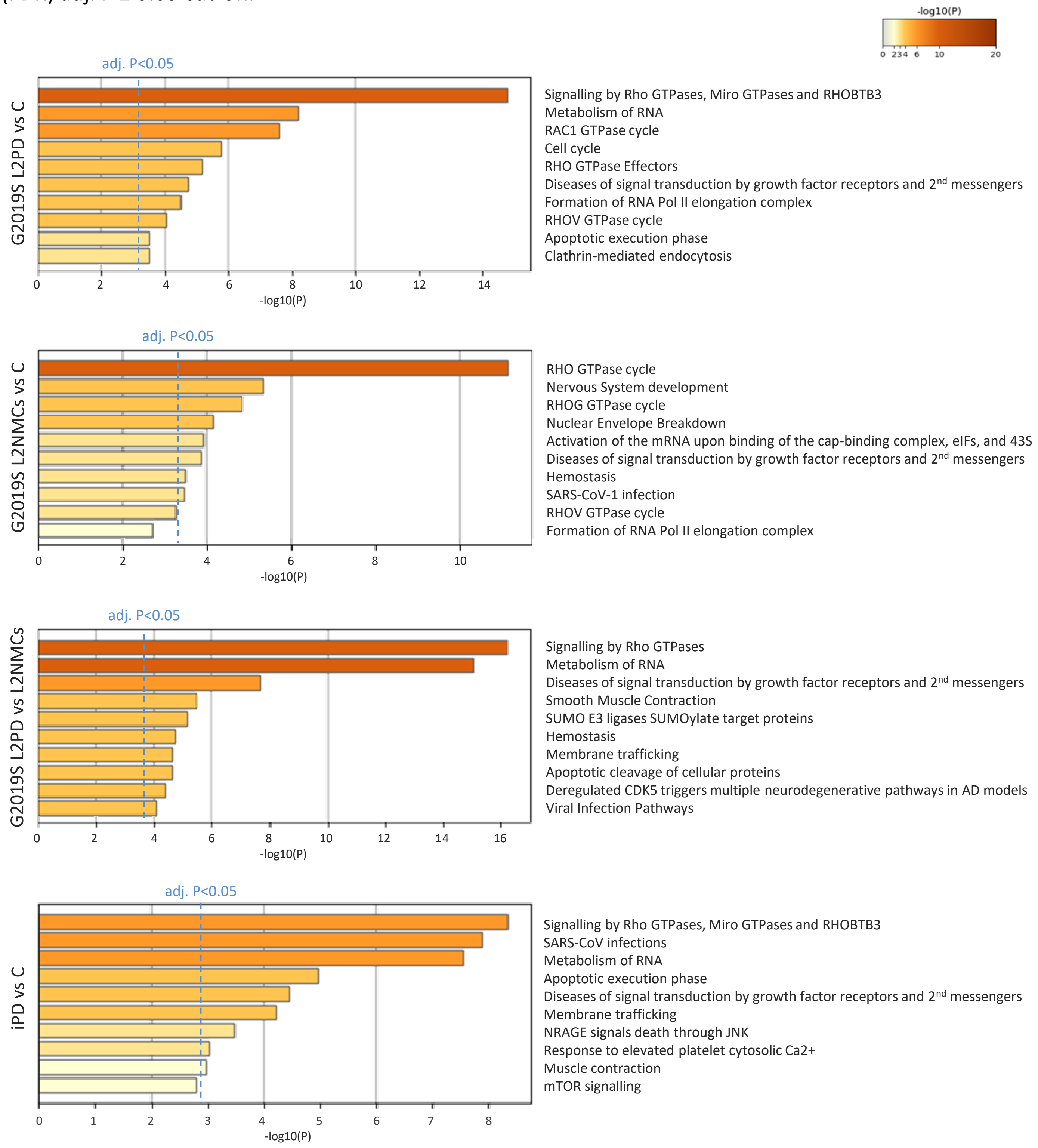

**Supplementary Figure 4. Immunoblot assessment of pThr37/46 4E-BP1 and total 4E-BP1 protein levels.** Full immunoblot assessment of pThr37/46 4E-BP1 and total 4E-BP1 using a subset of the cohort (n=24), including G2019S L2PD (n=6), G2019S L2NMCs (n=6), iPD (n=6), and controls (n=6).

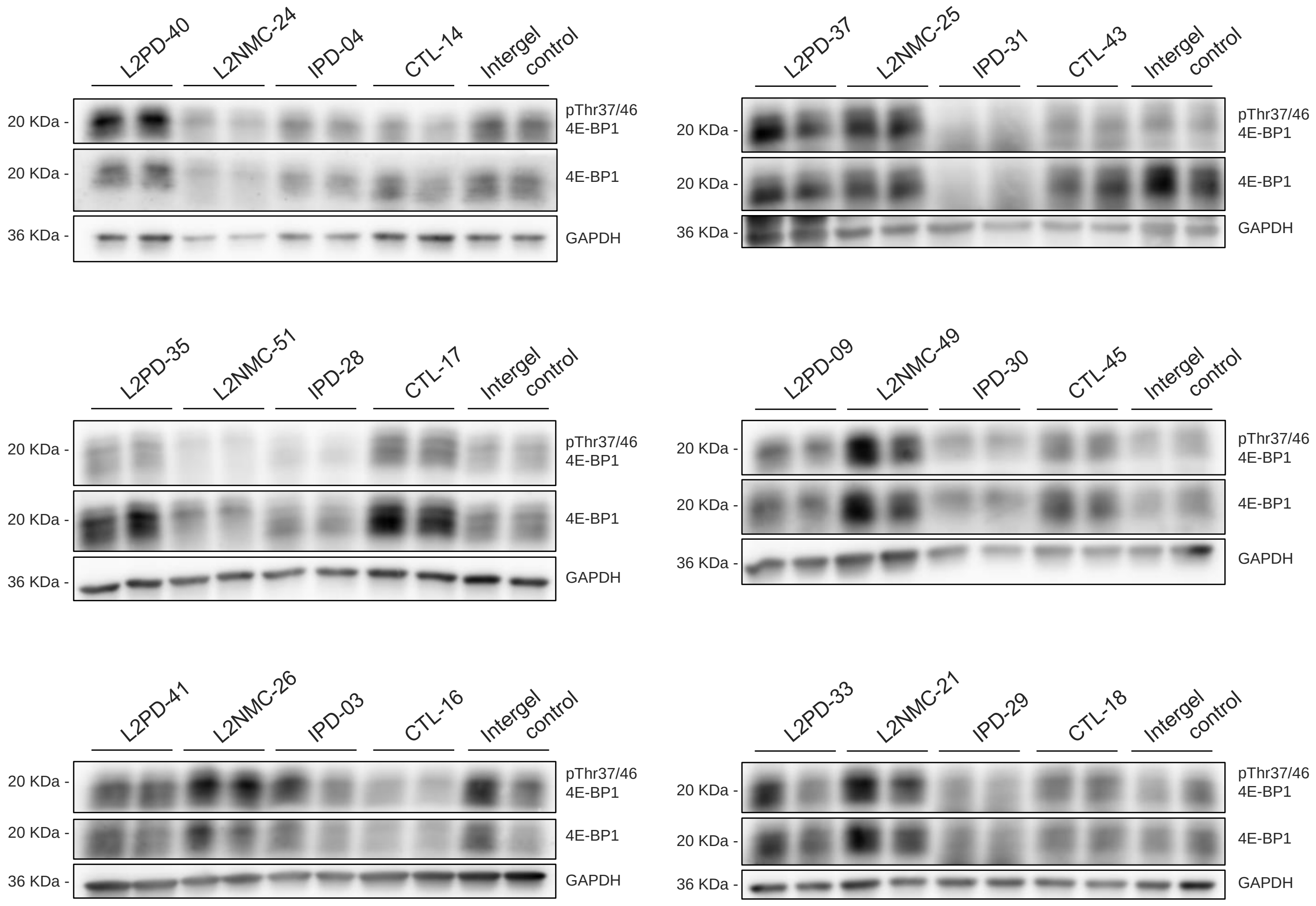

**Supplementary Figure 5. Immunoblot assessment of ATG9a and MRPS14 protein levels.** Full immunoblot assessment of ATG9a and MRSP14 using a subset of the cohort (n=24), including G2019S L2PD (n=6), G2019S L2NMCs (n=6), iPD (n=6), and controls (n=6). Boxplots representing normalised levels after the band densitometric analysis for the various studied makers in all subjects studied in duplicates, as follows: ATG9a / MRPS14 / GADPH, all of them double normalised to the same intergel control, also measured in duplicates.

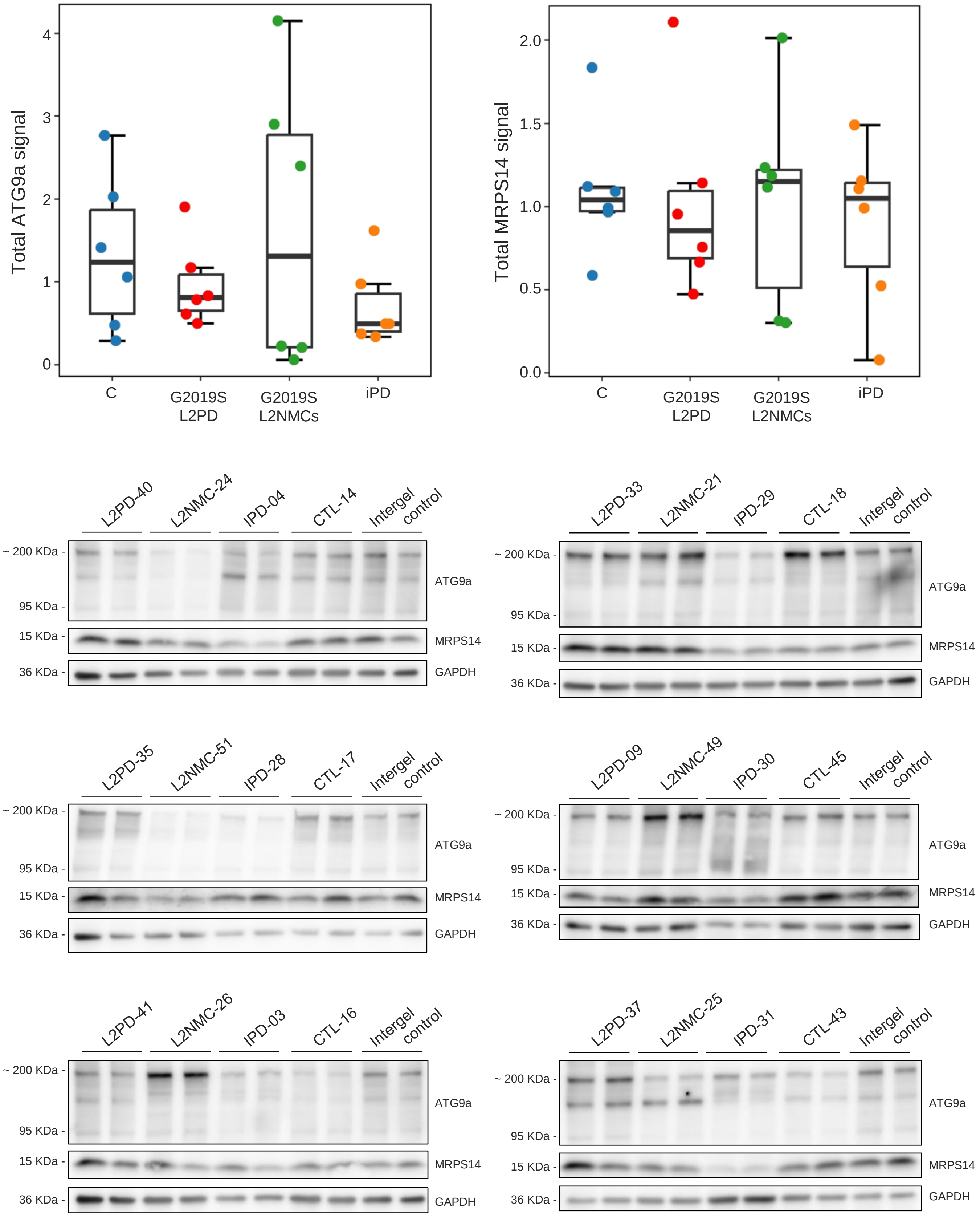

**Supplementary Figure 6 (i). Expanded pThr37/46 4E-BP1 and pSer106 RAB12 responsiveness to MLi-2 LRRK2 inhibition.** Full immunoblot analysis of pThr37/46 4E-BP1 / total 4E-BP1 and pSer106 RAB12 / total RAB12 using two technical replicates of fibroblasts from G2019S L2PD (n=4), G2019S L2NMCs (n=2), iPD (n=2) and healthy controls (n=2), treated with DMSO or the MLi-2 LRRK2 inhibitor.

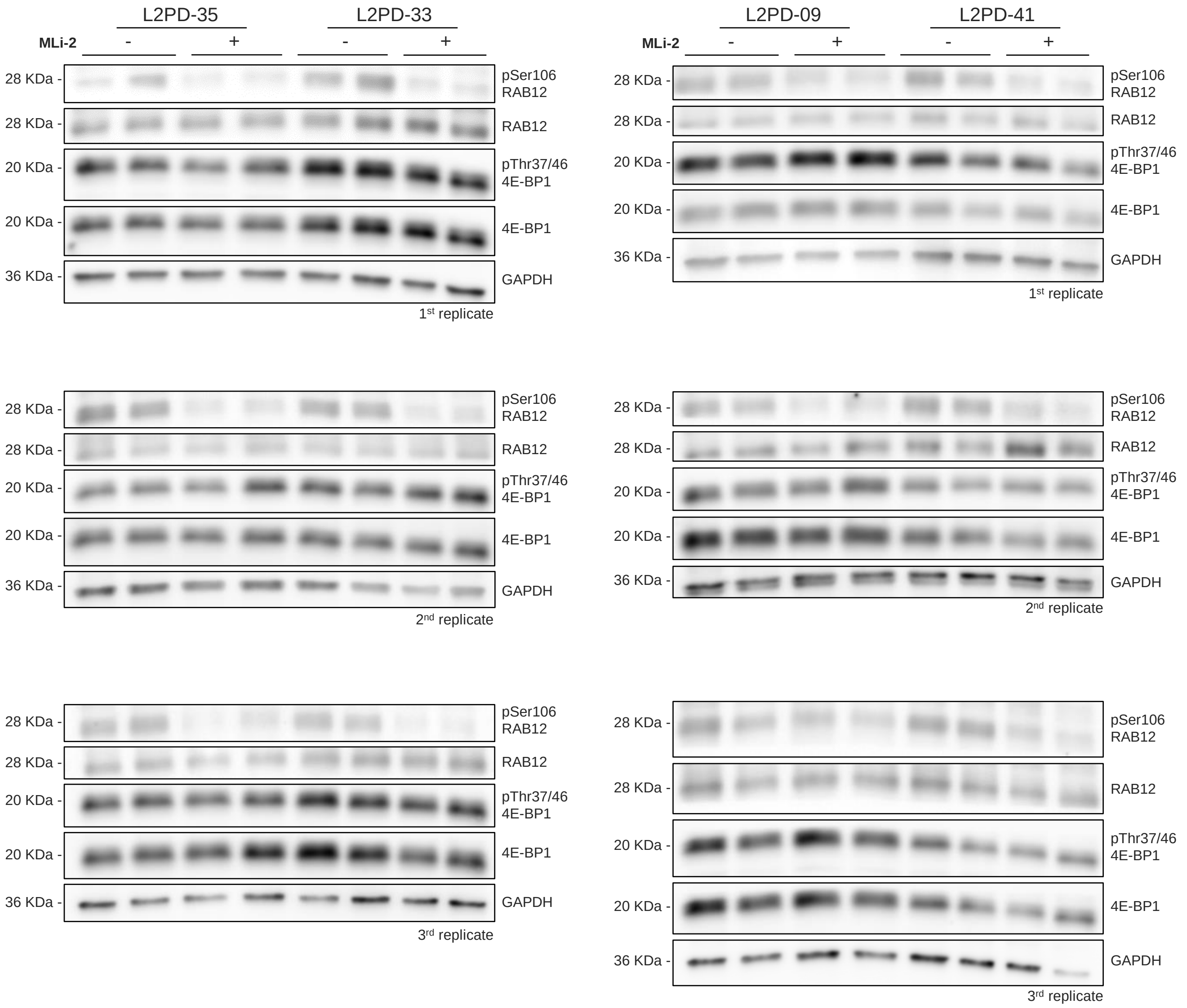

**Supplementary Figure 6 (ii). Expanded pThr37/46 4E-BP1 and pSer106 RAB12 responsiveness to MLi-2 LRRK2 inhibition.** Full immunoblot analysis of pThr37/46 4E-BP1 / total 4E-BP1 and pSer106 RAB12 / total RAB12 using two technical replicates of fibroblasts from G2019S L2PD (n=4), G2019S L2NMCs (n=2), iPD (n=2) and healthy controls (n=2), treated with DMSO or the MLi-2 LRRK2 inhibitor.

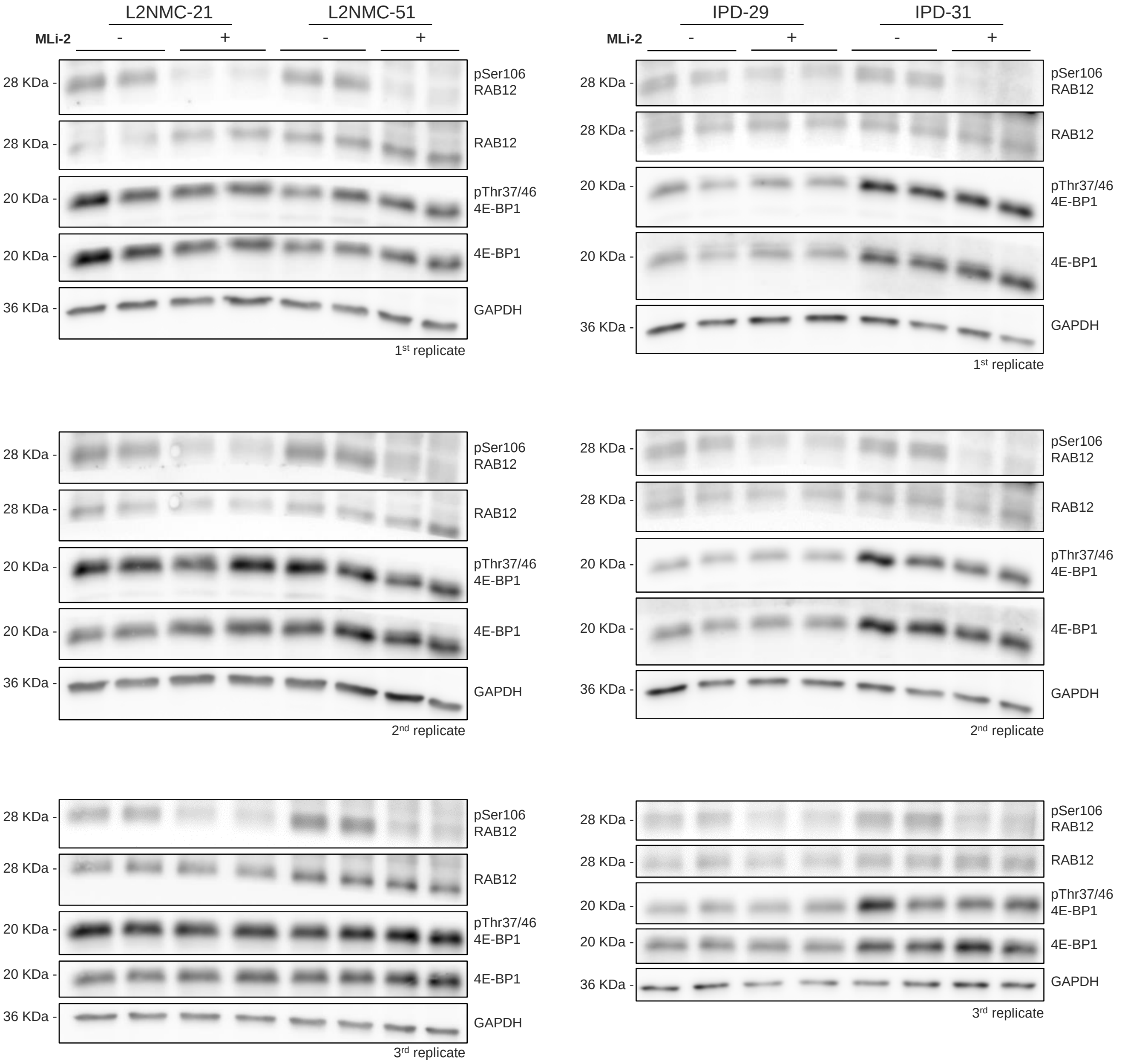

**Supplementary Figure 6 (iii). Expanded pThr37/46 4E-BP1 and pSer106 RAB12 responsiveness to MLi-2 LRRK2 inhibition.** Full immunoblot analysis of pThr37/46 4E-BP1 / total 4E-BP1 and pSer106 RAB12 / total RAB12 using two technical replicates of fibroblasts from G2019S L2PD (n=4), G2019S L2NMCs (n=2), iPD (n=2) and healthy controls (n=2), treated with DMSO or the MLi-2 LRRK2 inhibitor.

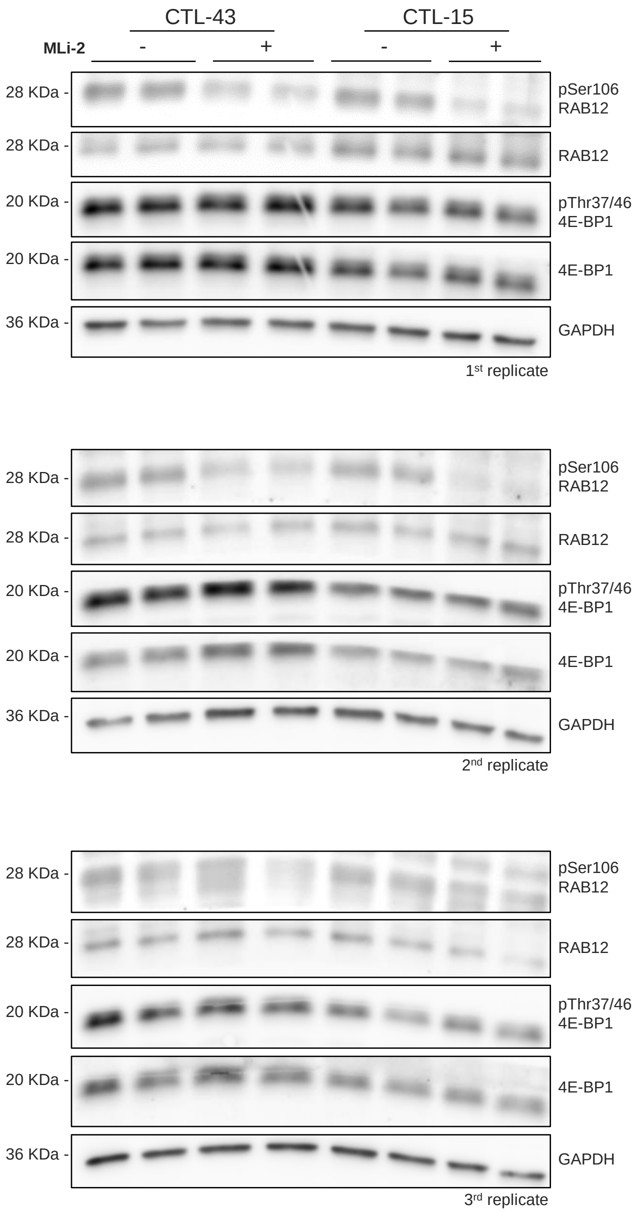

**Supplementary Figure 7. Expanded pThr73 RAB10 and pSer935 LRRK2 responsiveness to MLi-2 LRRK2 inhibition.** Full immunoblot analysis of pThr73 RAB10 / total RAB10 and pSer935 LRRK2 / total LRRK2 using two technical replicates of fibroblasts from G2019S L2PD (n=4), G2019S L2NMCs (n=2), iPD (n=2) and healthy controls (n=2), treated with DMSO or the MLi-2 LRRK2 inhibitor (200 nM, 30 min), showing a diminishment of pThr73 RAB10 and pSer935 LRRK2 phosphorylation levels after LRRK2 inhibition by MLi-2 treatment. * (P-value ≤ 0.05) ** (P-value ≤ 0.01) *** (P-value ≤ 0.001).

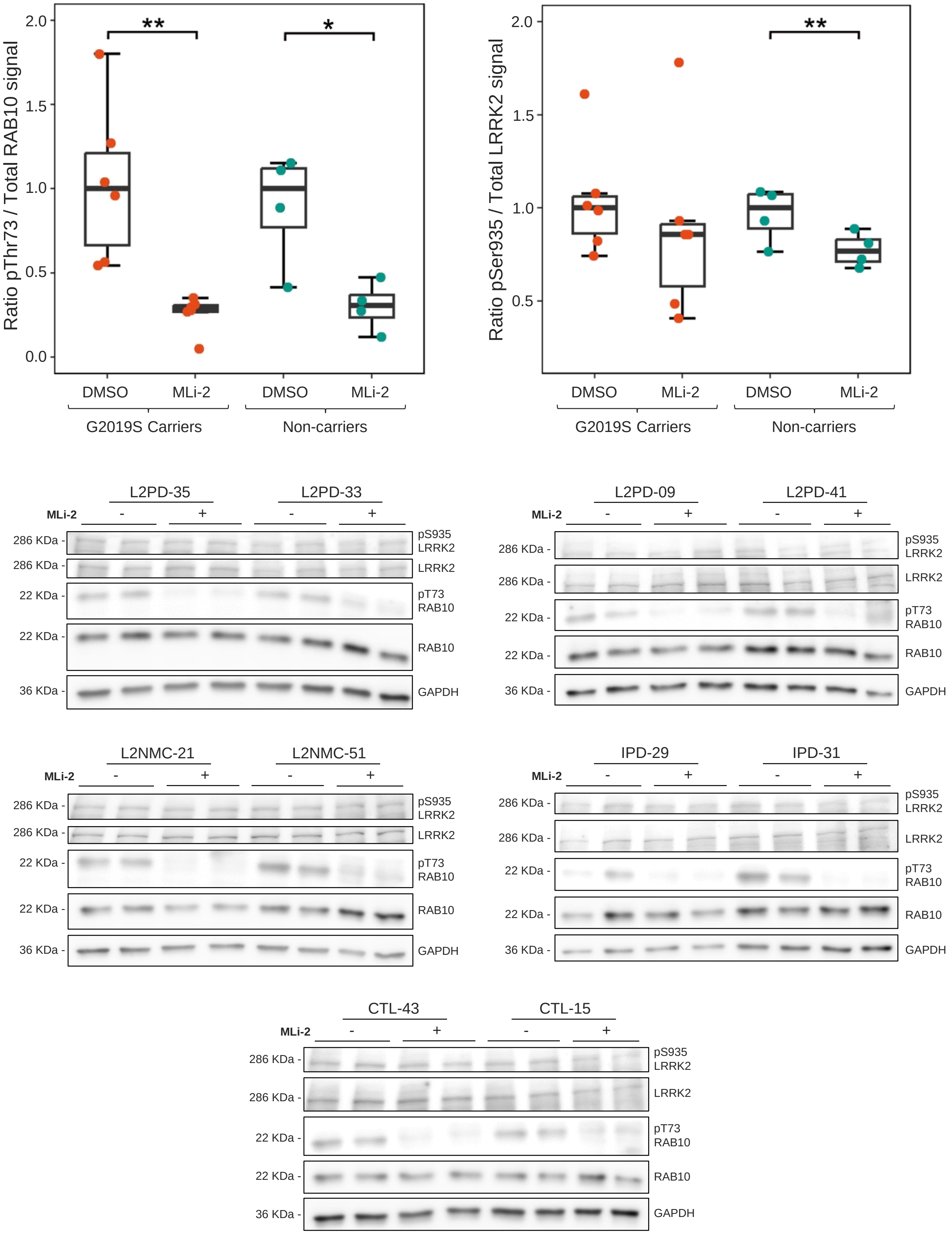
